# Neurofilament Light Disordered Tail Mutations Reshape Its Self-Assembled Network Structure

**DOI:** 10.64898/2026.03.27.714705

**Authors:** Rawan Aodeh, Yoav Dan, Dean Yona, Mohammad Shalabi, Asaf Sivan, Mathar Kravicas, Hillel Aharoni, Gil Koren, Lihi Adler-Abramovich, Roy Beck

## Abstract

Proteins with intrinsically disordered regions (IDRs) perform essential cellular functions despite lacking stable structures, challenging the traditional structure-function paradigm. Neurofilament-light (NFL) proteins assemble into bottlebrush filaments, whose disordered tail domains mediate nematic hydrogel formation critical for neuronal integrity. Mutations in NFL are linked to Charcot–Marie–Tooth (CMT) disease, yet their molecular effects remain unclear. Here, aiming to gain insight into these molecular mechanisms, we combine small-angle X-ray scattering, microscopy, and deep-learning conformational analysis to investigate CMT-associated NFL tail mutations. We find that these mutations induce pathological hydrogel compaction, disrupt filament nematic order by generating microdomains, and alter water retention dynamics by reshaping of sequence-dependent conformational ensembles, leading to macroscopic network rearrangements. These findings provide mechanistic insight into how subtle sequence changes in IDRs modulate protein network organization and function, informing an understanding of IDR-related pathologies and mutation-based disease characterization.

## Introduction

One of the most significant achievements in structural biology has been the establishment of the structure-function paradigm, linking the amino acid sequence of a protein with its three-dimensional structure and biological function^1–4^. This paradigm has successfully explained diseases caused by point mutations and has significantly advanced targeted drug design strategies^5^. However, the structure-function paradigm is challenged by functional intrinsically-disordered proteins and regions (IDP/Rs)^6^, which lack stable three-dimensional structures. IDP/Rs exhibit high flexibility, naturally adopting multiple conformations that facilitate diverse interactions essential for various cellular functions^1,6,7^. Importantly, IDP/Rs are not a niche domain, as recent studies suggest that approximately 50–70% of the human proteome includes functional IDRs that play critical roles in numerous biological processes and diseases^8^.

Inspired by polymer science, experimental, theoretical, and computational approaches have focused on characterizing and predicting the ensemble properties of IDP/Rs. Building on the success of deep-learning methods for structured proteins, similar strategies have been applied to predict macroscopic properties of IDRs^9^, such as their ensemble dimensions^10^ or interactions with neighboring macromolecules^11^. As with structured proteins, the amino acid sequence and specific patterns fundamentally govern the conformational ensembles of IDRs. For instance, charged and hydrophilic residues promote disorder, and the patterning of charges strongly influences the average expansion of IDP ensembles^12,13^.

IDP/Rs often show low sequence conservation because they lack the structural constraints that typically drive sequence preservation; however, in some cases, the presence of disorder itself is evolutionarily conserved even when sequence similarity is low^14^. Furthermore, since IDP/Rs share many properties with traditional polymers, the importance of specific molecular details or small sequence changes is often overlooked when considering their function. As a result, understanding how point mutations in IDRs can lead to disease phenotypes remains a major unsolved challenge^15^. In this study, we address this challenge by investigating the molecular consequences of such mutations.

Neurofilament (NFs) proteins contain long, flexible IDRs that fluctuate between multiple structural conformations^16^. NFs are the major cytoskeletal components of neurons, serving as mechanical shock absorbers that protect axons from mechanical stress^17^. NFs self-assemble into 10 nm filaments via coil-coiled interaction of the ordered rod domain, while the C-terminal disordered tail domain results in a distinctive brush-like architecture^18–24^ (Figs. 1a, b). The primary NFs subunit protein that homopolymerizes to form mature filaments is neurofilament light (NFL)^18^.

**Figure 1.**
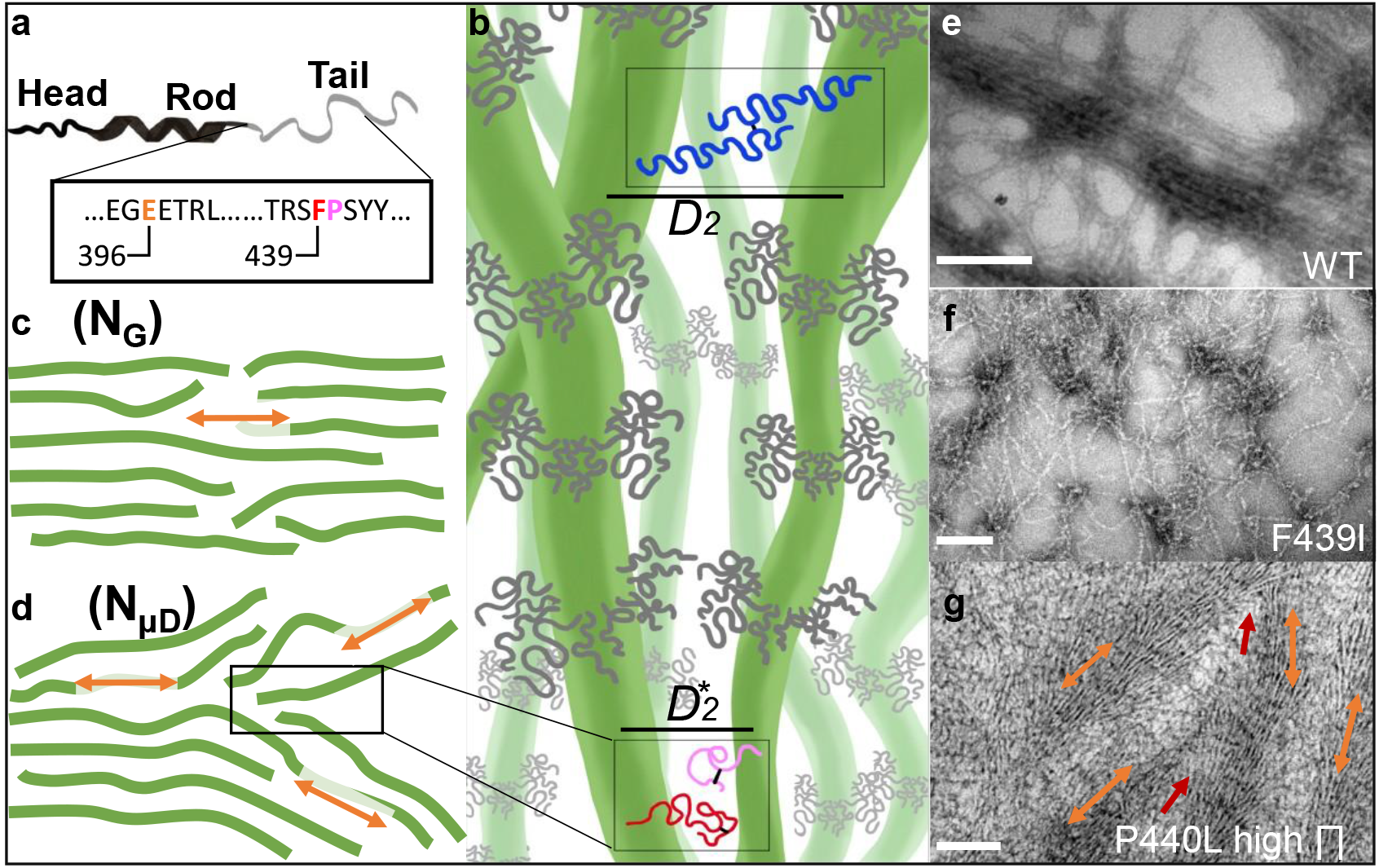
Self-assembly of NFL and effect of mutation on network architecture. **(a)** Schematic representation of the human NFL protein monomer, comprising an intrinsically disordered N-terminal head, a central folded helical rod, and a disordered C-terminal tail with pathological CMT-associated mutations highlighted. (b) The subunit proteins self-assemble into 10 nm-wide bottlebrush filaments via the rod domain, represented here as solid green rods. Inter-filament interactions are mediated by the disordered C-terminal tails projecting from the filament surface to form a bottlebrush architecture. Two cases are depicted: a greater filament spacing (D_2_), representing tail interactions at the expansion state, and a reduced spacing (D_2_*), demonstrating stronger tail interactions and a potential tail collapsed state. (c, d) At the larger mesoscale network level, two structural states are depicted: (c) nematic gel (N_G_) and (d) nematic microdomain (N_μD_). In the N_G_ phase, the NFL aligns into extended domains. In the N_μD_ phase, local defects separate bundled, aligned filament domains, creating water-rich gaps. Orange arrows indicate the local orientation of the nematic director. **(e-g)** TEM images of NFL and mutant samples. **(e)** WT NFL **(f)** CMT-mutant sample F439I measured without osmolytes and **(g)** P440L sample with added osmolytes (25wt% PEG) showing condensed and aligned filaments (orange arrows) with frequent nematic defects (red arrow). Scale bars, 200 nm.

Within myelinated axons, NFs are organized as parallel bottle-brush filaments arranged in a nematic liquid crystal phase. This nematic alignment is characterized by long-range orientational order and is driven by frequent interaction between neighboring C-terminal tails^25^. This nematic phase is fundamental to the biological function and the structural integrity of the neuronal cytoskeleton^25,26^. *In vitro* studies have shown that NF subunits (e.g., NF-L, NF-Medium, and NF-Heavy) can form distinct hydrogel mesophases, with their organization influenced by factors such as ionic strength and post-translational modifications (PTMs), most notably phosphorylation. While the heavier subunits (NF-M and NF-H) possess extensive phosphorylation repeats, the NF-L subunit contains one experimentally measured phosphorylation site in the tail domain^16,17,25,27–29^.

Genetic alterations in NF subunits are linked to Charcot–Marie-Tooth (CMT) disease^30^, the most common inherited neuromuscular disorder^31^. CMT is characterized by progressive, length-dependent degeneration of peripheral nerves^32^. Approximately 1% of diagnosed CMT cases are attributed to mutations in the NFL, with 8/30/7 mutations identified in the head/rod/tail domains, respectively^33,34^. Additionally, further mutations in these domains have been reported as pathological or likely pathological in clinical genomic studies^35^. Notably, previous reports have shown that several NFL mutants retain the ability to form filaments when co-expressed with other subunits ^36^.

Herein, we employ advanced analytical, X-ray scattering, and imaging techniques to investigate reconstituted human NFL protein, which self-assembles into hydrogel structures. Specifically, we evaluate the structural consequences of the CMT-related mutation in the NFL tail disordered domain that mediate interfilament interaction. We show that NFL tail mutants self-assemble into ∼10 nm-wide filaments while exhibiting altered macroscopic network structure, mechanical properties, and water-retention capabilities. Our experimental findings demonstrate that CMT tail domain mutations give rise to nematic microdomains, initiated by frequent, inherent defects in the long-range nematic order.

## Results

### Mutations Preserve Filament Self-assembly but Alter Network Organization

Similar to other intermediate filaments, native NFL protein self-assembles *in vitro* to form ∼10 nm wide bottle-brush filaments, which form a stable hydrogel at high filament density^37^. Here, we investigated the effect of point mutations at the carboxy-terminal region of the IDR in recombinant human NFL, which are not expected to affect filament formation^38^ (Fig. 1a). The CMT-linked mutations studied are E396K (located at the last residue of the rod domain), F439I, and P440L. Additionally, we included three control mutations, F439I/P440L, F402I, and P470L. The first double mutation variant aims to evaluate any effects of cooperativity between the mutations, while the latter two, neither of which has been associated with clinical pathology, involve the same amino-acid substitutions as in the CMT-related mutations but at different positions. A list of all studied variants is given in Table 1.

**Table 1.** Structural and mechanical properties of NFL hydrogels at *C*_*s*_=160 mM.

| | FW<br>(nm) | LC<br>domains | $\phi_D$ map | A map | CPM<br>color | $\chi$ plot | $\Pi_2$<br>(kPa) | $\Pi_\mu$<br>(kPa) | $D_0$<br>(nm) | $T_D$<br>(hr) | $T_R$<br>(hr) |
| --- | --- | --- | --- | --- | --- | --- | --- | --- | --- | --- | --- |
| WT | 10.2<br>$\pm 1.98$ | Strong<br>large<br>domains | Ordered | High<br>(red)<br>Low<br>(blue) | Uniform | High | 26.6 | 26<br>7 | 52 ±<br>5.9 | 2.5 | 4.6 |
| F439I | 10.04<br>$\pm 2$ | Strong<br>large<br>variation | Hetero-<br>geneous | High<br>local | Variable | Low | w/o | w/<br>o | 47 ±<br>0.9 | 3.3 | 6.3 |
| P440L | 13.86<br>$\pm 1.78$ | Strong<br>large<br>domains | Domain-Dominant | Uniform | Uniform | None | 107 | 53<br>5 | 36 ±<br>0.6 | 4 | 6.3 |
| E396K | 11.95<br>$\pm 2.27$ | Weak,<br>micro-<br>domains | Hetero-<br>geneous<br>(blue) | high<br>(red) | Variable<br>(blue) | None | w/o | w/<br>o | 55 ±<br>2.9 | 3.3 | 5.3 |
| F439I/P440L | 11.03<br>$\pm 1.8$ | Strong<br>large<br>domains | Domain-Dominant | Uniform | Uniform | High | w/o | 15<br>39 | 37 ±<br>1.4 | 3.3 | >6.3 |
| F402I | 11.85<br>$\pm 2.27$ | Strong<br>large<br>domains | Domain-Dominant | High<br>(red) | Uniform | High | 26.6 | 26<br>7 | 49.5<br>$\pm 3$ | 3.6 | 3.1 |
| P470L | 12.99<br>$\pm 2.66$ | Weak<br>micro-<br>domains | Hetero-<br>geneous<br>(blue) | High<br>(red) | Variable<br>(blue) | Low | w/o | w/<br>o | 37 ±<br>0.4 | 3.6 | >6.3 |

Initially, transmission electron microscopy (TEM) confirmed that the mutant variants self-assembled *in vitro* into long, flexible filaments ∼10 nm in width, similar to the wild type (WT) protein (Figs. 1e-f, S2)^27^. In all samples, the filaments’ length was much larger than their width. At higher filament densities, WT sample exhibited a more uniform and aligned filament organization, whereas the mutant samples tended to form branched bundle topologies (Figs. 1b-g).

### Mutations Affect Macroscale Filament Alignment and Induce Nematic Microdomains

At high filament density, native NFs condense into an aligned hydrogel network through interactions between tail-tail IDRs^16,17,25,27,28^. To evaluate the effects of mutations on the macroscopic alignment of the NF network, we employed cross-polarized microscopy (CPM) and synchrotron small-angle X-ray scattering (SAXS).

Under near-physiological salt conditions, all large NFL hydrogels exhibited birefringent domains (Figs. 2a-d, S4). However, we identified two distinct macroscopic organizational phases; (a) in WT, F402I hydrogels, as well as in the P440L and F439I/P440L pathological mutants, we observed an extended nematic gel (N_G_) mesophase characterized by macroscopic aligned filaments and large birefringent domains (Fig. 2c); and (b) in the F439I, and E396K hydrogel, as well as in the P470L control, we observed an extended nematic microdomains (N_μD_) mesophase, where filament alignment was broken into smaller microscopic domains with localized orientation, visualized as colorful birefringent regions under CPM (Fig. 2d).

**Figure 2.**
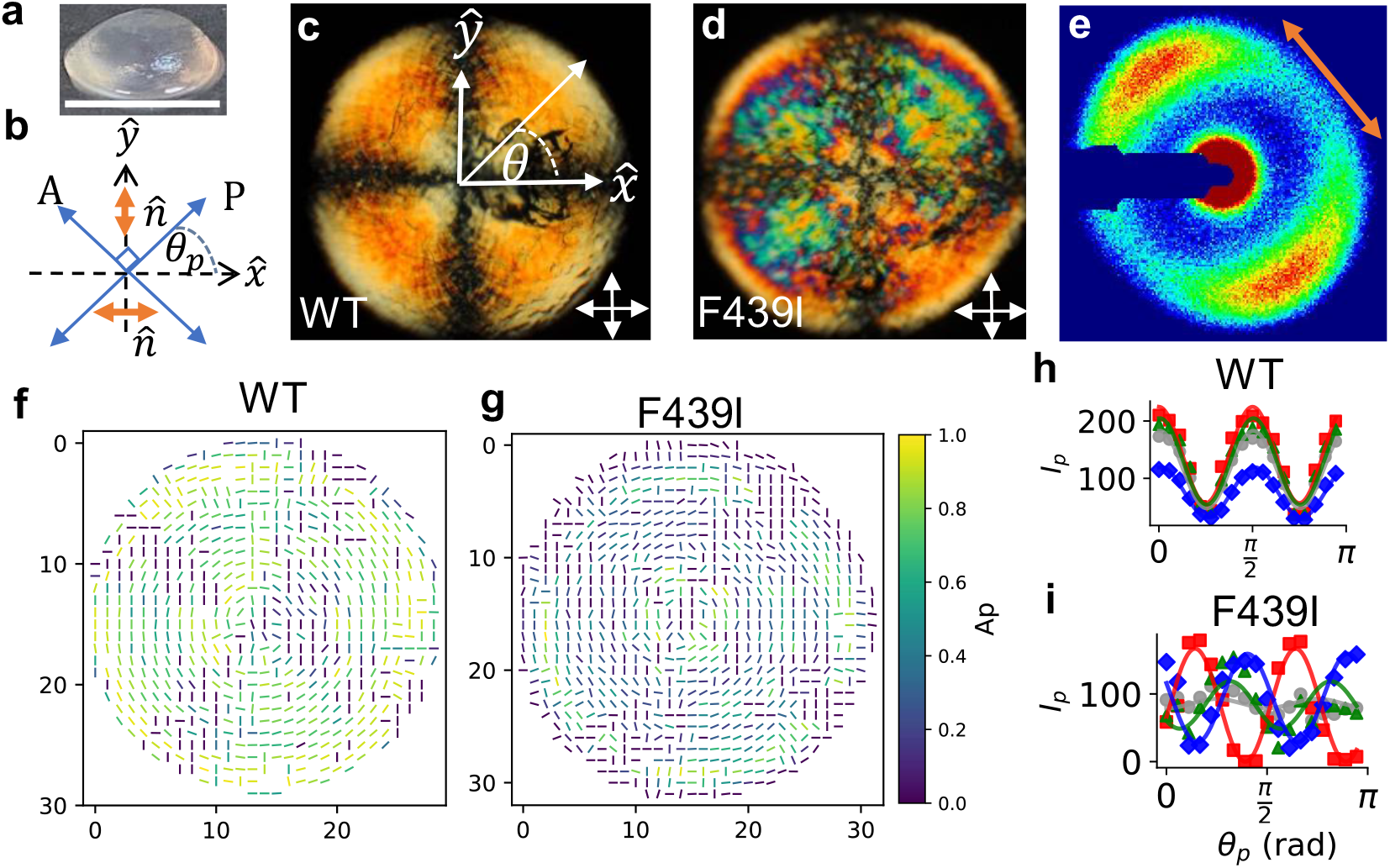
Macroscale filament order in NFL hydrogels assessed using CPM and scanning SAXS. **(a)** Large NFL hydrogel sample (approximately 7.5 mm in diameter, and 1.5 mm in height) exhibiting a lens-like shape that was prepared under near-physiological salt concentration (*C*_*s*_ = 160 mM). **(b)** Schematic illustrating the CPM geometry. The axes of the polarizer (P) and analyzer (A) are indicated by blue arrows, with both elements rotated by an angle *θ*_*P*_ relative to the sample. For nematic orientation along the x or y axes (orange arrows), the maximal CPM intensity will be when θ_P_=π/4+nπ/2, where n is an integer number. **(c, d)** Representative CPM images of WT and F439I mutant hydrogels. WT displays uniform birefringent regions with nematic alignment, featuring an S = +1 defect at the center. F439I exhibits pronounced N_μD_ with prominent reddish and bluish regions. **(e)**, Representative anisotropic 2D SAXS scattering pattern indicating local filament orientation (orange arrow). **(f, g)** CPM director direction maps were generated by extracting local director orientation using Equation 1. The intrinsic π/2 phase ambiguity was relieved by identifying the director orientations using scanning SAXS and Equation 2 (Fig. S4) **(f)** WT exhibits highly ordered, symmetrically patterned phase distributions indicative of robust long-range nematic order, whereas **(g)** F439I shows pronounced spatial heterogeneity characteristic of N_μD_ formation. **(h, i)** Quantitative intensity profiles from selected hydrogel regions (Fig. S3). Data is shown for grayscale, red, green, and blue channels. (**h)** WT demonstrates consistent high-intensity maxima across all color channels, while **(i)** F439I shows substantial differences in the director orientation among the different color channels.

To spatially analyze the nematic local director orientation (*φ*_*D*_), we captured CPM images at different polarizer angles relative to the sample (*θ*_*p*_), while keeping the analyzer cross-polarized (Fig. 2b-d, supplementary movies). Here, the local light intensity 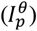 can be fitted to:

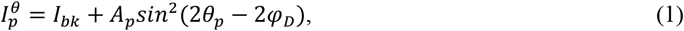

where *θ*_*p*_ is the angle between the sample and the polarizer, and *φ*_*D*_ is the director’s orientation, *A*_*p*_ is the modulation amplitude, and *I*_*bk*_ is the background intensity. The modulation amplitude *A*_*p*_ extracted from this fitting quantifies the degree of local nematic alignment. Importantly, when 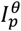 is maximized, the filaments are either parallel or perpendicular to the bisection of the polarizer and analyzer angles (i.e., at *φ*_*D*_ ±π/4, see Fig. 2b). However, because the CPM technique cannot distinguish between filament orientations differing by π/2, there is an intrinsic phase ambiguity.

To resolve this limitation, we identified the local filament direction of the CPM using synchrotron scanning SAXS data. Here, to extract local anisotropy, the 2D data were integrated radially surrounding the correlation peak position (q=0.087-0.29 nm^-1^) and plotted against the azimuthal scattering angle (χ), as shown in (Figs. 2e and S3). Empirically, we found the χ-plots fit:

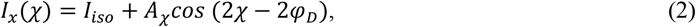

here *A*_χ_ represents the anisotropy amplitude, *φ*_*D*_ indicates the preferred director orientation in real space, and *I*_*iso*_ is the isotropic background. Therefore, for each anisotropic position measured by CPM, we could resolve the phase ambiguity (*φ*_*D*_ or *φ*_*D*_+ π/2) by selecting the orientation that best matched the SAXS-derived direction. These orientations served as focal points for neighbors with lower anisotropy, whose orientations were subsequently considered. The orientations selected were those that most closely resembled a continuous orientation gradient (Figs. 2f, g and S4).

Consistently, all mutant samples exhibited more local defects compared to the WT sample (Fig. S4). Furthermore, the WT, F439I, E396K, F402I, and P470L hydrogels exhibited a central bend-dominated +1 topological defect with planar anchoring at the sample periphery. Notably, the anisotropy signal was markedly reduced both at the sample center due to the loss of nematic order at the central defect, and at the periphery, due to reduced sample thickness from the lens-like geometry of the hydrogel (Figs. 2f, g, and S4). In contrast to the WT, almost no anisotropy profiles were detected for P440L and F439I/P440L in the scanning SAXS experiments (Fig. S4),. The scanning SAXS data reveals scale-dependent structural alignment, where the P440L and F439I/P440L mutants exhibit broad birefringent regions, suggesting weak nematic alignment, also demonstrated under qualitative CPM analysis (Fig. 3a). Moreover, the nearly isotropic SAXS profiles for P440L and F439I/P440L indicates that its ordered nematic domains are highly localized with typical domain sizes are smaller than the X-ray beam size (∼200×90 µm^2^).

**Figure 3.**
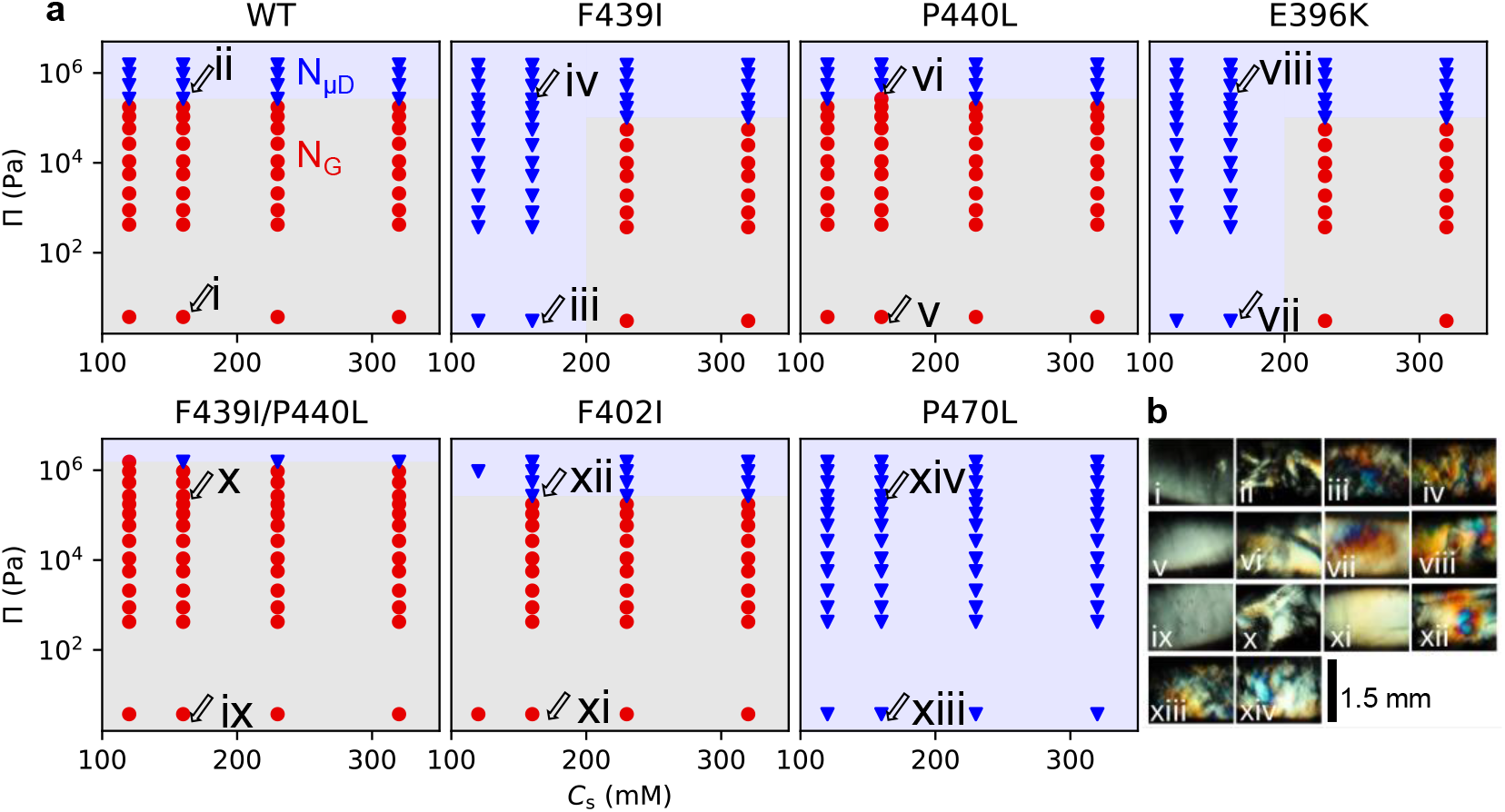
Phase diagram and CPM images of NFL hydrogel. **(a)** Osmotic pressure (*Π*) versus (*C*_*s*_) phase diagram, showing conditions under which NFL networks form nematic gel (N_G_, red circle) or nematic microdomains (N_μD_, blue triangle), as determined by CPM. **(b)** Representative CPM images of hydrogels prepared in quartz capillaries (1.5mm width). At low *Π* (without PEG), WT networks form extended nematic alignment, whereas CMT-related mutants (F439I, E396K), and P470L control mutation already display N_μD_. At high *Π* (<15wt% PEG), all samples form N_μD_ characterized by birefringent microdomains with distinct color variation. The phase diagram shows that mutations shift the boundary between the N_G_ and N_μD_ phase, with mutants exhibiting disrupted nematic order at significantly lower *Π* and Cs than WT. The specific CPM images for the samples are marked accordingly in panel **a**.

WT, P440L, F439I/P440L, and F402I hydrogels exhibited homogeneous birefringence across all CPM color channels (Figs. 2h, S5, S6), with WT samples showing the most intense signal. In contrast, the CPM for the F439I mutant showed colorful microdomains, substantial variation in *φ*_*D*_ across the color channels, and heterogeneity in local director orientation (Figs. 2d, g, i). E396K and P470L mutants displayed intermediate behavior, with some variation in *φ*_*D*_ across the color channels, particularly in the blue channel (Figs. S5 and S6). These results demonstrate that NFL tail mutation disrupts the network’s global order, creating fractured ordered domains.

CMT (F439I, P440L and E396K) and control variants (F439I/P440L, F402I and P470L) are highlighted in red and green, respectively. Filament width (*FW*): extracted from TEM images; Liquid Crystal (LC) domains: specify the size and degree of filament alignment under low *Π*, quantified from SAXS azimuthal scans and CPM images; Director orientation (*φ*_*D*_) and amplitude (*A*) maps: specify the spatial heterogeneities as measured from SAXS scans and CPM; CPM color: indicates local director agreement among the different color channels. Specific color indicates the channel with a different *φ*_*D*_-value from the other colors; *χ*-plot: indicates the level of anisotropy found in the SAXS-scan 2D images; *Π*_*2*_: is the lowest osmotic pressure at which two SAXS correlation peaks first appear, marking the onset of a new structural arrangement; *Π*_*μ*_: is the lowest osmotic pressure at which CPM detects the N_μD_ phase transition by small colorful domains. w/o indicates appearance of N_μD_ without any PEG additions; *T*_*D*_ and *T*_*R*_: are the times for the hydrogel to lose/regain 50 % of its initial weight under dehydration/rehydration conditions, respectively.

### Mutants Tune NFL Network Structural Properties

NF-NF interactions are mediated by transient ionic cross-bridges that can be tuned by buffer salinity. These interactions, in turn, are responsible for network stiffness. Following, we evaluated the modulation of the macroscopic alignment of WT and mutant networks under various ionic strengths (Cs=120-320 mM) and osmotic pressures (*Π* = 1 - 10^6^ Pa). Specifically, *Π* was modulated by varying polyethylene glycol (PEG) concentration in the buffer solution^39^.

Here, smaller hydrogel samples were inserted into quartz capillaries, imaged under CPM, and then measured using SAXS. Phase transitions in these capillary-based experiments were identified from the macroscopic optical appearance of the hydrogels, using birefringent intensity and colorfulness as primary indicators for phase assignment. The specific geometry of these sample holders (1.5 mm quartz capillaries), optimized for standard SAXS data collection, prevents quantitative spatial mapping of the local director. Instead, these measurements here were focused on characterizing the global macroscopic phase behavior across a wide range of ionic strengths and osmotic pressures.

Notably, all samples showed colorful N_μD_ phase at high *Π* (Fig. 3, blue triangles). However, CMT-related mutations (F439I and E396K) exhibited a N_μD_ phase even at low *Π* (0% w/v PEG) and low ionic strengths (120 and 160 mM) (Fig. S7). In contrast, the P440L and F439I/P440L mutations resulted in nematic domains across a large salinity and *Π* regime (Fig. S7).

The specific position of the mutation within the disordered tail influenced the extent of nematic organization. The F402I control mutation resulted in nematic alignment similar to WT across a broad salinity range, whereas the P470L control mutant exhibited a more pronounced and persistent N_μD_ phase compared to both WT and P440L under similar conditions (Fig. S7). These observations are consistent with previous reports indicating that inter-filament interactions are primarily governed by the peripheral extensions of the disordered tail domains^17^.

Using SAXS, we investigated the effect of mutations on the interactions between neighboring filaments. Here, the inter-filament distance, *D*, is probed from the nematic correlation SAXS peak positions (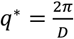, see Fig. S8). Uncharacteristic of polymeric brushes, we found that even a single point mutation in the long-disordered tail domain significantly modulated the inter-filament distance, *D* (Fig. 4). For example, at *C*_*s*_ = 230mM, the WT and the E396K, and F402I mutations exhibited an expanded network with larger osmolyte-free spacing *D*_0_ ≈ 60 nm (Figs. 4b, d). In contrast, the F439I, P440L, and P470L mutations formed more condensed networks with a smaller *D*_0_ ≈ 50 nm. The F439I/P440L double mutant sample exhibited a cumulative effect, resulting in the smallest osmolyte-free spacing *D*_0_ ≈ 40 nm (Figs 4b, d).

**Figure 4.**
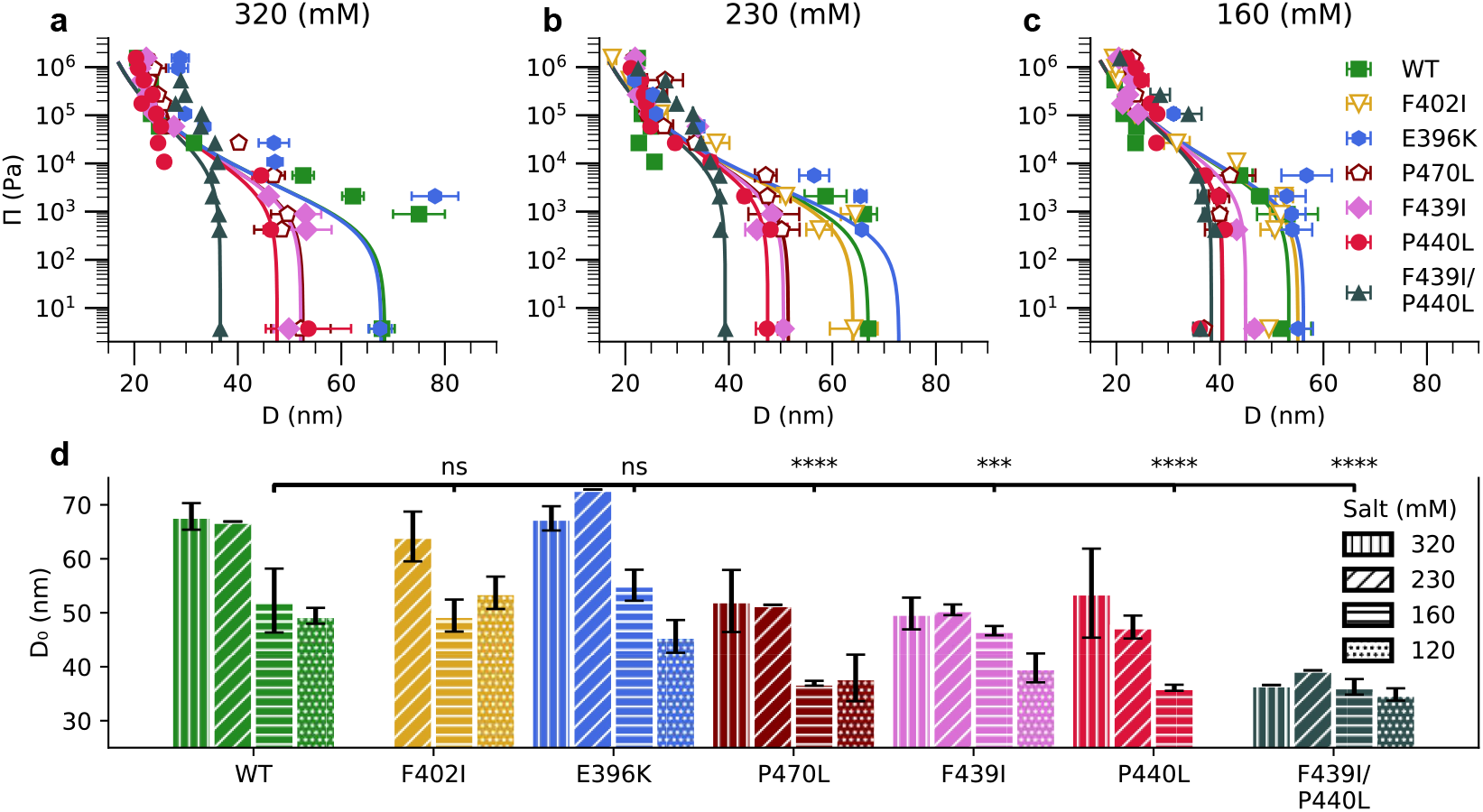
Osmotic compression and local conformational states of NFL hydrogels. **(a-c)** Osmotic pressure (*Π)* versus interfilament spacing (D) for WT NFL, CMT-linked mutation (E396K, F439I, P440L, and F439I/P440L - filled symbols) and control mutant (F402I, P470L - empty symbols) samples at various salt concentrations (*C*_*s*_ = 320, 230, 160 mM) measured by SAXS. The solid lines fit a mean-field theory of hydrogel compression^17^. Error bars represent sample heterogeneity arising from measurements taken at different positions along the hydrogel. The data at *C*_*s*_ =120 mM deviate significantly from the model^17^. **(d)** Dunnett’s multiple comparison test was performed to determine if the uncompressed spacing (*D*_*0*_) significantly differed from the WT sample at *C*_*s*_ = 160 mM. F402I and E396K were not significantly different (ns), while mutants P440L, F439I/P440L, and P470L showed extremely significant differences (****, p < 0.0001), and the F439I mutant showed very highly significant differences (***, p < 0.001).

At higher *Π*, all NFL hydrogels transitioned into highly condensed states, reaching *D*_1_∼20 nm. As previously reported for bovine NFL, at low salinity (120 mM), the hydrogels’ ability to condense under high *Π* was significantly impacted, and the scattering peak was broadened ^17,27^. The *Π*-*D* response can be fitted to the theoretical mean-field model^17^ using a single fitting parameter, k, representative of the number and strength of the attractive interaction domains between the IDR tails of adjacent filaments (Figs. 4d and S9).

Moreover, all samples exhibiting colorful N_μD_ mesophase under CPM showed a secondary SAXS correlation peak. In fact, at the lowest osmotic pressure, we found a second correlation peak (*Π* ≡ *Π*_2_) preceding the one we identified with N_μD_ using CPM (*Π*_*μ*_ ≥ *Π*_2_, table 1). The only exception was the F439I/P440L double mutant, where *Π*_*μ*_ ≫ *Π*_2_ = 0, for which the network was much denser in the absence of added osmolytes. The second peak is indicative of an alternative larger structural ordering (Figs. S8e, f) with highly densely packed filament bundles (with interfilament distance *D*_1_) that are separated by larger gaps (*D*_2_, see Fig. 1). Note that the second peak did not exhibit any clear trend with variations in *Π*, and its value remained consistently large (*D*_2_ ∼ 80 − 100 nm, Fig. S10 black line). Moreover, the secondary peak was more pronounced in the SAXS scans on the larger hydrogel samples, clearly showing microdomain boundaries (Fig. S11a).

### CMT-linked Mutation Alters the Local Ensemble Conformations

To assess whether CMT-related mutations alter the local ensemble conformation, we synthesized 23-residue long peptides corresponding to the WT and the F439I, P440L and F439I/P440L mutants spanning positions 434-456 (Fig. 5a). Using flow-cell SAXS, both WT and F439I showed typical polymeric patterns of IDP with a continuous rise in the Kratky representation at higher q (Iq^2^ vs. q, Fig. 5b), while, notably, the F439I peptide exhibited a higher plateau compared to the WT. Using extended Guinier analysis^40^ (Figs. S12a, b), the WT and F439I peptides exhibited a radius of gyration 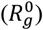 of 1.46±0.02 and 1.42±0.01 nm, and Flory scaling exponent (*ν*^0^) of 0.604±0.001 and 0.606±0.002, respectively (Table S1). These small differences are consistent with disorder analysis showing that the F439I mutation is predicted to have structural disorder similar to the WT and different from the P440L variant (Figs. S13a-c).

**Figure 5.**
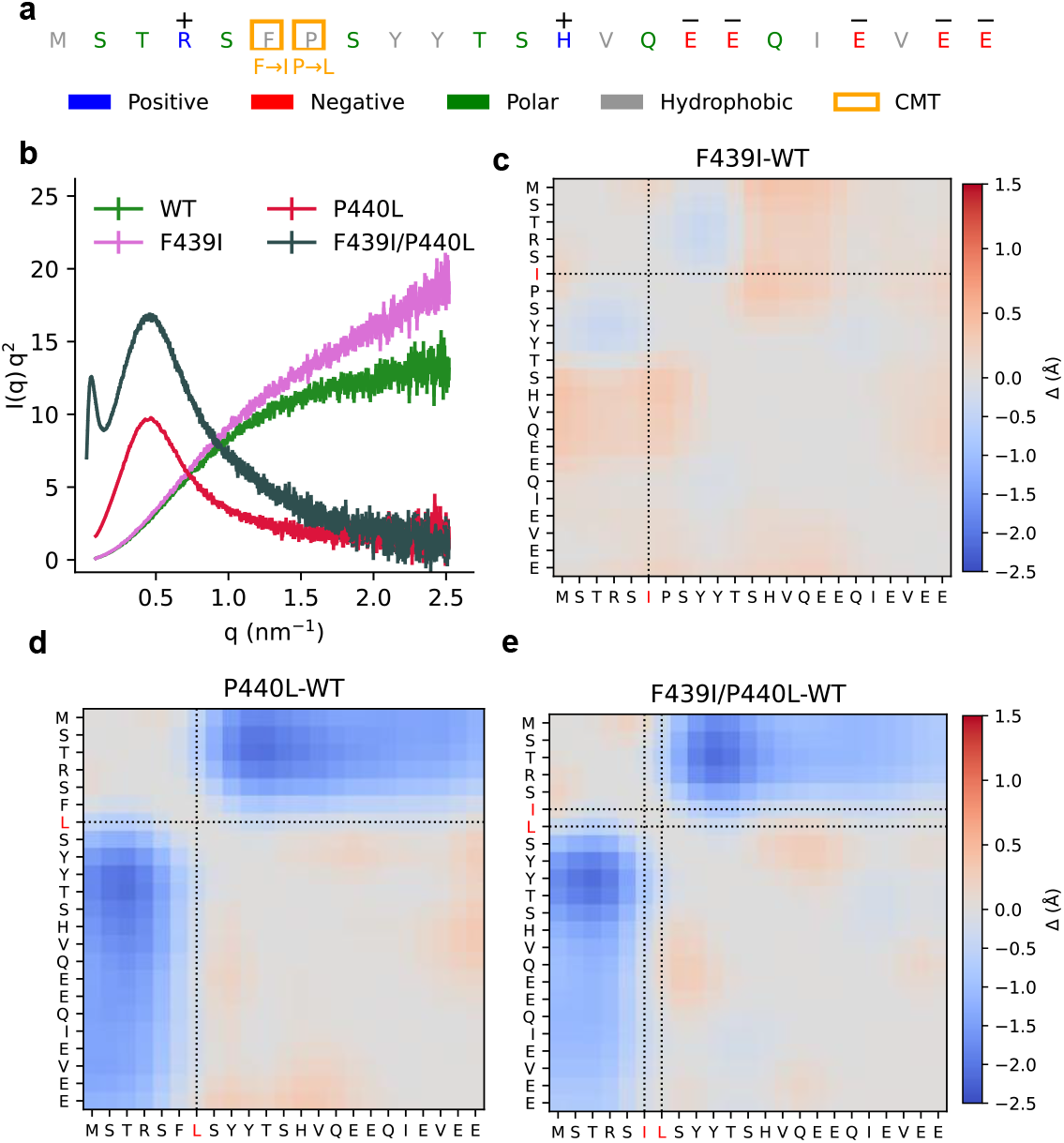
Short peptide conformational states. **(a)** Sequence of 23-residue NFL tail peptide, highlighting amino acid properties: positive (blue), negative (red), polar (green), hydrophobic (gray), and residues associated with CMT-linked mutations (orange). **(b)** Kratky plots (Iq^2^ vs. q), comparing the conformational properties and flexibility of WT, F439I, P440L and F439I/P440L peptides. WT and F439I peptides exhibited scattering profiles typical of IDPs. In contrast, P440L and F439I/P440L samples aggregated rapidly upon solvation, displaying a characteristic bell-shaped Kratky profile with pronounced low-*q* scattering, indicative of aggregation and the formation of compact, multimeric globular structures. **(c-e)** Deep-learning STARLING^41^ prediction of residue-residue distance maps subtracted from the WT prediction (Δ). Predication variation for sequences of the 23mer peptides were experimentally measured using solution-SAXS (F439I, P440L, F439I/P440L). In agreement with the experiments showing aggregation, P440L and F439I/P440L sequences show post-mutation reduced residue-residue distances.

Nonetheless, the P440L and F439I/P440L mutant peptides aggregated immediately upon solvation in the physiological salinity buffer. The compacted globular structure was further validated by SAXS, as evidenced by a bell-shaped Kratky plot (Fig. 5b). Here, we noted that the F439I/P440L mutant displayed a distinct dual-peak feature in the Kratky plot, suggesting the formation of heterogeneous, multi-scale-regulated aggregates (Fig. 5b).

To disrupt intermolecular aggregation, the peptides were measured at elevated urea concentrations (Table S1). Under these denaturing conditions, the WT and F439I peptides retained IDP profiles, while the P440L and F439I/P440L peptides continued to display a compact, aggregated conformation, evident from increased scattering at low *q*-values. Due to the aggregation behavior of P440L and F439I/P440L, Guinier analysis was not applicable. Instead, the SAXS intensity at zero angle, *I*_*0*_, was extrapolated (Fig. S12c, d) to estimate the average aggregation number of the interacting peptides (Table S1). Surprisingly, increasing the urea concentration to as much as 2M lead to higher aggregation numbers for both peptides (Table S1)

To support our mutation-induced structural ensemble results, we employed STARLING^41^, a recently deposited deep learning algorithm, for predicting conformational ensemble contact maps of IDRs (Fig. 5c-e). In agreement with the peptide’s aggregation experiments, compared to the WT, STARLING predicted nearly unchanged contact maps for F439I (Fig. 5c) but smaller average intermolecular distances pockets, *i*.*e*., compaction, in the vicinity of the P440L, or F439I/P440L mutations. Moreover, when employed across the entire disordered tail, STARLING showed local compaction in the vicinity of the mutations and expansion to residues farther away (Fig. S14).

### Hydrogel Water Retention is Mutant Dependent

Defects in the nematic order can generate hydrated gaps between filament-rich and aligned domains that can influence water retention capabilities^25^. Given the correlation between point mutations and microdomain mesophase formation (Figs. S4 and S7), we examined the weight loss/gain of large samples under identical hydration/dehydration conditions. Weight loss and gain were normalized to the initial (*m*_0_ ≈ 65.2 ± 9.45 mg) and final (*m*_*d*_ ≈ 8.24 ± 2 mg) dehydrated weights.

The WT hydrogels released and absorbed water in minimal time (Figs. 6a, b). The other N_G_ samples, P440L and F402I, were slowest to dehydrate but were similar to the WT in rehydration dynamics (Figs. 6a, b). The F439I, E396K, and P470L N_μD_ samples, as well as the dense F439I/P440L N_G_ sample, showed intermediate behavior. Despite the variation in kinetics, none of the hydrogels returned to their original weight after 48 hours, consistent with irreversible compaction under high *Π*, as previously reported ^28^.

**Figure 6.**
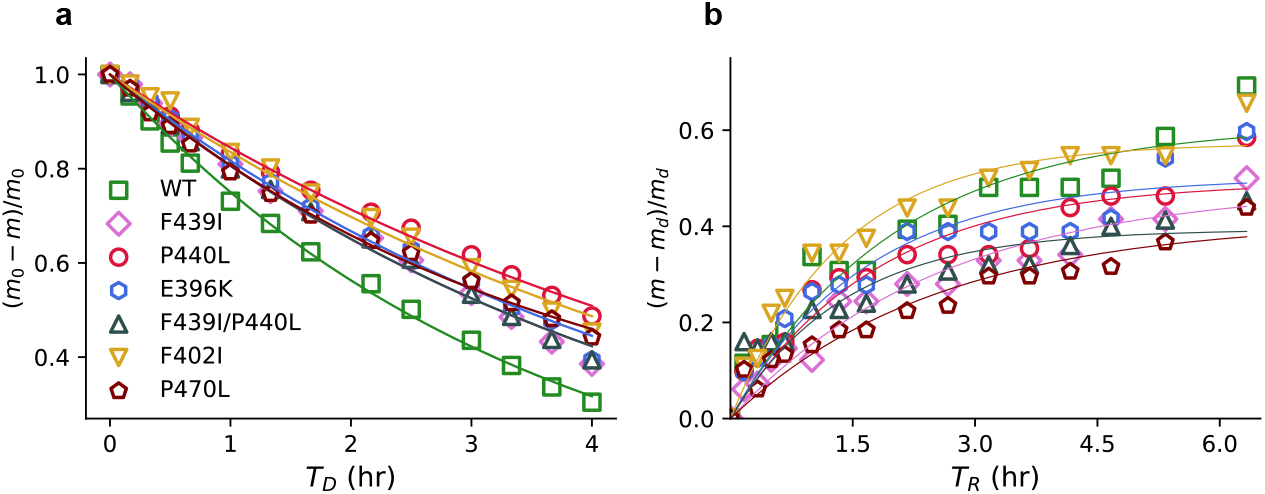
Hydration dynamics characterization of NFL mutant hydrogels. **(a)** Spontaneous dehydration, and **(b)** rehydration profiles. WT hydrogels exhibited the fastest water loss and gain, while the mutants displayed variable kinetics. P440L and F402I showed the slowest dehydration, and F439I, F439I/P440L, and P470L showed moderate dehydration rates and slow rehydration. Curves were fitted using an asymptotic regression model. Together, the results show that mutations not only alter nematic order (N_G_ vs N_μD_) but also modify water-retention properties, with disrupted nematic alignment leading to slower hydration dynamics.

## Discussion

Functional analysis of mutations in folded proteins has been a cornerstone in structural biology and medicine. However, due to the structural plasticity, low sequence homology, and polymeric statistical nature in IDP/Rs, it is natural to assume that point mutations that preserve amino acid properties will bear a less significant structural and functional effect. Our study challenges this assumption by demonstrating that a point mutation in the NFL disordered tail domain can result in substantial structural and functional alterations.

The functional role of the NFL disordered tail domains is to transiently crosslink neighboring filaments and to maintain long-range nematic order, which is directly correlated to the NF-network structure^17^. TEM imaging revealed intact filament formation, accompanied by altered larger-scale organization, in the tail mutant variants. This result is in agreement with previous studies, proposing that CMT might not result from a failure of NF to self-assemble into filaments but rather from a disorganization of the NF along axons, leading to focal accumulations^36^. It further supports studies showing that nematic long-range order and regulated inter-filament spacing are critical for supporting neuronal integrity and function^37,42^. Therefore, the studied point-mutation modulations of local order, mechanical, and hydration properties (Table 1) are consistent with the associated disease pathology^43^.

Our findings indicate that most point mutations led to a more compact, stiffer hydrogel network. The exceptions are F402I and E396K, where the mutations are located closer to the bottlebrush backbone. This result is consistent with previous studies showing that network spacing and stiffness are correlated with inter-filament interactions, which are expected to be most significant towards the tips of the tails^17,27^. This spatial hierarchy explains why E396K and F439I display different nanoscopic spacing despite both forming fragmented macroscopic phases (Fig.3). Because distal tail extensions primarily mediate inter-filament distance, the backbone-proximal E396K mutation preserves WT-like internal spacing within its microdomains. Conversely, the distal F439I and P440L mutations directly alter these peripheral interactions, driving local pathological compaction. Consequently, while both proximal and distal mutations disrupt long-range macroscopic nematic order, SAXS captures their distinct nanoscopic architectures: expanded WT-like spacing for E396K, and condensed spacing for F439I.

Effective tail length can be reduced by increased intramolecular attractions that loop the disordered tails onto themselves or toward the hydrophobic backbone. Supporting loop formation is demonstrated by the short-peptide experiments where P440L and F439I/P440L mutants showed dominant aggregation propensity (Fig. 5b). These results demonstrate enhanced intramolecular attractions for these tails, which is also supported by the predicted contact maps using the deep-learning STARLING algorithm^41^ (Fig. 5c-e). Interestingly, these specific mutations induce a highly stable, collapse-prone conformational state that resists chemical denaturation. Nonetheless, increased attractions likely promote local tail “looping,” effectively shortening the disordered tail and reducing its repulsive volume. This behavior provides a direct molecular link to the pathologically condensed inter-filament spacing observed in the full-length mutant hydrogels (Fig. 4).

Supporting our results is the recent proline–leucine (PL) repeat simulation revealing that hydration properties in IDP/Rs are highly sequence-dependent^44^. In particular, clusters of hydrophobic residues, such as leucine, have been shown to enhance hydrophobic interactions, resulting in local structural compaction^44^. Indeed, replacing proline with leucine (P440L) notably increased hydrophobic-driven aggregation and peptide chain compaction (Figs. 5b and S12). This transition is further mediated by the loss of P440 unique cis-trans isomerization^45^, which provides a structural mechanism to regulate the local conformational ensemble. Eliminating this conformational barrier facilitates the rapid peptide collapse observed in our experiments, leading to misfolding and aggregation. These residue-specific insights illustrate how minor amino acid substitutions significantly influence structural dynamics and illuminate sequence-dependent factors that govern the behavior of IDP/Rs.

An alternative means for a denser network is through the disorder-order transition. Sequence-based bioinformatic predictions suggest the studied mutations may influence local disorder within the tail domain (Fig. S13). Notably, F439I was predicted to increase disorder, while E396K and P440L were associated with a more ordered state. These predictions are consistent, although with different magnitudes, across multiple prediction tools. The control mutations, such as F402I and P470L, exhibited similar trends but with more modest effects.

The transient interaction between neighboring filaments, mainly through ionic bridging between tails, is functionally critical for supporting the mechanical properties of neurofilament hydrogels^16,25,28^. Previous studies on NF hydrogels showed a narrow salinity range between the nematic and isotropic mesophases, where the sample transitions to a bluish-opaque gel (B_G_) phase with microphase separation^25^. While B_G_ appeared in non-physiological salinities, the macroscopic organization resembles the N_μD_ we report here. The weak secondary SAXS correlation peaks (Figs. S10 and S11) strongly indicate the presence of competing interfilament spacing with similar free energies. At the network level, these alternative free-energy minima lead to excess topological defects (Figs. 1 and 7). This explains the observed correlation between microphase separation, as measured by CPM (Figs. S4 and S7), and the secondary SAXS correlation peaks (Figs. S10 and S11). At high osmotic pressures, where the disordered tails are further compressed, the N_μD_ phase becomes dominant across all samples. However, the specific mutation determines the condition (crowding or salinity) under which such defects are more likely to occur. Previous studies showed that the native NF network’s nano- and macroscopic structures depend on PTMs^16,46,47^. We therefore expect that mutations that induce structural modifications may be coupled to these PTMs, particularly for the longer subunit proteins (i.e., NFM, NFH) that have abundant phosphorylation sites.

**Figure 7.**
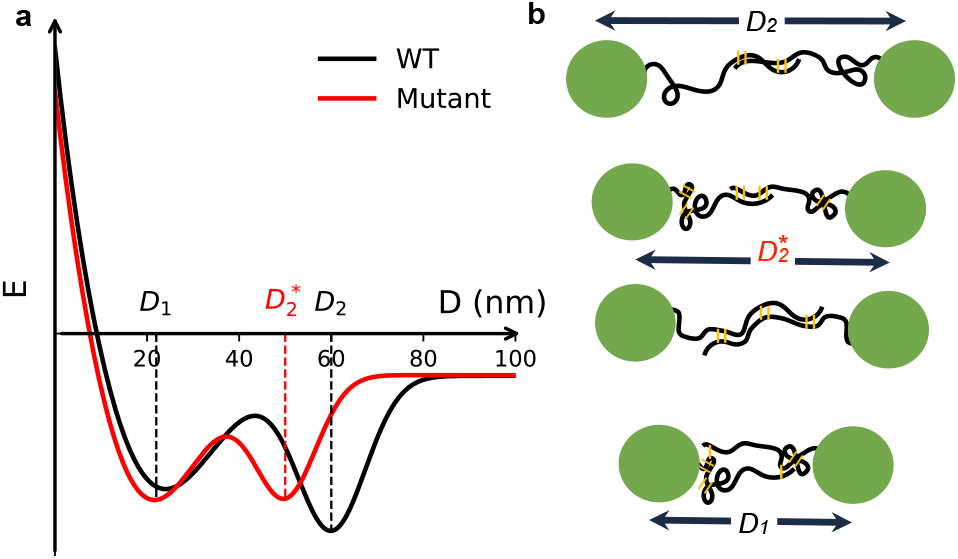
Schematic representation of the NFL network energy landscape and possible tail conformations. Free energy profile vs. inter-filament spacing (D) schematics for WT and mutated NFL network. WT protein networks exhibit an expanded state (D = D_2_) minimum, and an energy barrier to a secondary condensed state (D = D_1_) that becomes accessible at high osmotic pressure. The network of mutant proteins shows an energy minimum for the expanded state at smaller inter-filament spacing (D = D_2_*) and a shallower energy barrier to the condensed state, as evident from the two SAXS correlation peaks (Fig. S8) of the coexisting states. **(b)** Schematic illustration of NFL tails conformations and crosslinks (yellow lines) at different states. WT proteins (top) show extended tail conformations with crosslinking mainly at the tail’s tips. Mutated proteins (middle) potentially result in local compactions (e.g., P440L, F439I/P440L) or additional overlap between neighboring tails (e.g., F439I). Highly entangled tails state (bottom) at high osmotic pressure, where they show large overlap and additional crosslinking.

The mutation-dependent water retention results can be explained by two competing factors. On the one hand, frequent local disruptions in network alignment (N_μD_ mesophase) induce larger water-rich reservoirs, slowing water release and absorption (e.g., F439I, E396K, P470L mutations). Similarly, slower hydration dynamics were observed in the B_G_ phase at lower salinities, where disruption of the nematic phase creates large water reservoirs^25^. On the other hand, the denser networks (e.g., P440L, F439I/P440L) are inherently dehydrated and thus water exchange between the filaments is limited. The structure-function relationship is further demonstrated by the WT and the F402I mutant, which show similar water retention properties, nanoscopic interfilament spacing compression profiles, and N_G_-N_μD_ phase diagrams (Figs. 3-5).

## Conclusion

In summary, our results establish a direct experimental link between single-residue mutations in the IDR tail of NFL and alterations in the biophysical network properties. By integrating comprehensive experimental data across multiple structural levels, we demonstrate that CMT-associated mutations in these IDRs do not affect individual filament assembly but alter the structural, mechanical, and hydration properties of NFL networks. Due to the relatively flat energy landscape of IDRs^48^, point mutations that maintain their physicochemical class (i.e., charge, hydrophobicity) can still introduce competing orders, as observed here.

Computational models for IDRs are still in their infancy and are largely based on simulations^8^. Nonetheless, as we demonstrate here, combining these models with experimental verification can yield valuable molecular insights. The findings presented here show a direct link between IDR sequence and ensemble properties. Other pathological point mutations in IDRs should provide a fruitful ground for similar investigations. It may also advance our understanding of CMT disease progression, highlighting the critical role of the tail IDRs in NFL protein interaction and network function. Further investigations should extend beyond the NFL alone, evaluating how these mutations interact with additional subunit proteins (e.g., NFM, NFH) that copolymerize with NFL, to fully elucidate the complexity of NF network dynamics in health and disease contexts. This will be addressed in future work.

## Supporting Information

The Supporting Information includes detailed Materials and Methods for NFL protein expression and purification, site-directed mutagenesis, synchrotron SAXS measurements, TEM imaging, and computational data analysis. It contains Figures S1–S14 and Table S1 as referred to in the main text and within the supporting experimental descriptions.

## Acknowledgments

Synchrotron SAXS data were collected at beamlines P12 (EMBL, DESY, Hamburg, Germany), I22 (Diamond Light Source, Didcot, UK), SWING (SOLEIL, France), and BM29 (ESRF, Grenoble, France). We thank Dmytro Soloviov (EMBL), Andy Smith (Diamond Light Source), Thomas Bizien (SOLEIL), Javier Perez (SOLEIL), and Mark Tully (ESRF) for their assistance during data collection. We thank the CCiTUB TEM-SEM Unit (Universitat de Barcelona), Itai Cabilly for 3D printing, and Asma Matar for graphic design. Special thanks to Dr. Anthony Brown (The Ohio State University) and Erwin Frey (LMU University) for critical reading and suggestions.

