## Supplemental File for "Neurofilament Light Disordered Tail Mutations Reshape Its Self-Assembled Network Structure"

This file contains:

- Material and methods
- Figures S1-S14
- Equations S1-S8
- Table S1

### Materials and Methods

### Experimental Section

#### Plasmid and Mutagenesis

The human wild-type (WT) NFL plasmid (phNFL, was a gift from Anthony Brown, Addgene plasmid #132589; http://n2t.net/addgene:132589; RRID)[1] served as the initial template. The NFL cDNA was amplified via PCR and cloned into a pET30 vector using NEBuilder HiFi DNA assembly. Site-directed mutagenesis PCR introduced CMT-associated mutations (F439I, P440L, E396K, F439I/P440L, F402I, and P470L) using the WT plasmid as the template. All mutations were confirmed through Sanger sequencing.

#### Protein Expression and Purification

Protein purification was performed as previously described[2]. Escherichia coli BL21(DE3) competent cells were transformed with the hNFL-pET construct and plated on agar supplemented with kanamycin. A single colony inoculated a 50 mL Luria-Bertani (LB) overnight culture with 50 µg/mL kanamycin at 37 C°. Subsequently, the overnight culture was scaled up into 1 L TB in baffled flasks and grown at 37 C°, 180 rpm, until reaching an optical density (OD600) of 0.6–1.0. Protein expression was induced by adding IPTG to a final concentration of 0.4 mM, followed by an additional 4-hour incubation. Bacteria were harvested via centrifugation and stored at −80 C°. For cell lysis, frozen pellets were resuspended in lysis buffer (50 mM Tris, pH 8; 100 mM NaCl; 1% Triton X-100), sonicated on ice (5-second pulses, 30% amplitude, total of 4 minutes), and centrifuged at 18,500g for 70 minutes at 4 C°. This sonication-centrifugation step was repeated twice. The resulting pellet was resuspended in a denaturing buffer (8 M urea, 100 mM sodium phosphate, pH 7) and centrifuged at 6,000g for 10 minutes. The supernatant was purified via DEAE Sepharose column chromatography using a gradient elution with 200 mM NaCl. The collected NFL fractions were concentrated and subjected to size-exclusion chromatography. Protein purity (>95%) was confirmed by SDS-PAGE gel (Fig. S1). Purified NFL proteins were concentrated and dialyzed into assembly buffer (100 mM MES, pH 6.8 adjusted with NaOH; 0.02% $NaN_3$; 1 mM EDTA; 7 mM β-mercaptoethanol). The total ionic strength was adjusted by varying NaCl concentrations to 120, 160, 230, or 320 mM. Assembly proceeded for 48 h at 37 °C.

#### Hydrogel Preparation

After dialysis/assembly, NFL filaments were pelleted by ultracentrifugation. For capillary SAXS/CPM**,** sample were spun at 50 000 rpm for 1 h (TLA-100 rotor), loaded into 1.5 mm quartz capillaries**,** and overlaid with PEG solutions to apply osmotic pressure (below) and sealed with epoxy glue to prevent dehydration. For large-hydrogel scanning SAXS/CPM, sample (sedimented with TLS-55 rotor) were mounted in a sealed mica-window holder prefilled with assembly buffer ($C_s$=160 mM) to prevent dehydration.

#### Osmotic Pressure Calibration

Capillaries were overlaid with approximately 100 µL of polyethylene glycol (PEG, 20,000 g/mol) at defined weight percentages and sealed to prevent dehydration. The corresponding osmotic pressures ($\Pi$) were determined using the PEG osmotic stress calibration according to [3]:

$$log_{10}^{\Pi} = 1.57 + 2.75(wt\%)^{0.21} \qquad \text{(S1)}$$

**Transmission Electron Microscopy (TEM)**

Diluted WT and mutant NFL protein samples were prepared by drop-casting 7 µL solutions onto 400 mesh copper grids and allowed to adsorb for 1 minute. Excess liquid was blotted away, followed by negative staining with 7 µL UranylLess solution (Bar Naor LTD) for 1 minute. Excess stain was again removed by blotting. Samples were imaged using a JEM 1400plus electron microscope operated at 80 kV and J1010 Transmission electron microscope (Jeol) coupled with an Orius CCD camera (Gatan) operated at 80kV. TEM imaging with the J1010 was carried out at the TEM-SEM Electron Microscopy Unit of the Scientific and Technological Centers (CCiTUB), Universitat de Barcelona.

**Cross-polarized optical microscopy (CPM)**

Capillary samples were imaged before SAXS measurement to validate the presence of the nematic liquid crystal phase. For larger hydrogel sample quantitative analysis, pixel intensities were fitted to:

$$I_p^{\theta} = I_{bk} + A_p sin^2(2\theta_p - 2\varphi_D) \tag{S2}$$

where $\theta_p$ is the angle between the sample and the polarizer, and $\varphi_D$ is the director's orientation, $A_p$ is the modulation amplitude, and $I_{bk}$ is the background intensity.

Importantly, when $I_p^{\theta}$ is maximized, the filaments are either parallel or perpendicular to the bisection of the polarizer and analyzer angles (i.e., at $\varphi \pm \pi/4$, see Fig. 2b). However, because the CPM technique cannot distinguish between filament orientations differing by $\frac{\pi}{2}$, there is an intrinsic phase ambiguity.

To resolve this limitation, we compared the filament direction from CPM to synchrotron SAXS data. For each anisotropic position, we considered two CPM candidates ($\varphi_D$ and $\varphi_D + \pi/2$) and selected the orientation that best matched the SAXS-derived direction. These orientations served as focal points for neighbors with lower anisotropy, whose orientations were subsequently considered. The orientations selected were those that most closely resembled a continuous orientation gradient.

**Small-angle X-ray scattering (SAXS): Data collection**

**Capillary hydrogels**: Preliminary experiments were performed at our home lab using an Eiger2R 1M detector and a Xenocs GeniX Low Divergence CuKα radiation source setup with scatterless slits. Subsequent measurements were performed at synchrotron facilities: beamline I22, Diamond Light Source, Didcot, UK; beamline P12, EMBL, DESY, Hamburg, Germany; and beamline SWING, SOLEIL, France.

**Larger NFL hydrogel:** Spatially resolved (scanning) SAXS maps were collected at beamline P12, EMBL, DESY, Hamburg, Germany. For scanning measurements, the hydrogel sample holder cavity was filled with assembly buffer ($Cs$ = 160 mM) to prevent dehydration, and the hydrogel dimensions were matched to the holder aperture to ensure the hydrogel remained securely in place. Between scans, the surrounding buffer was exchanged for solutions with higher PEG concentrations, and the hydrogels were allowed to equilibrate before additional measurement.

**Peptide solution:** Short 23-mer peptides (residues 434–456) were synthesized, verified by mass spectrometry, and purified to >95% by high-performance liquid chromatography (HPLC) by LifeTein LLC (Hillsborough, Nj, USA). Lyophilized powders were dissolved in assembly buffer containing 0, 1, 2, and 4 M urea. Next, we filtered and concentrated the samples using 3 kDa Ultra-centrifugal filters. Initial peptide concentrations were determined using a Nanodrop 2000 spectrophotometer (Thermo Scientific), where the maximum peptide concentrations reached were

5.0, 5.0, 4.0 and 4.0 mg/mL for WT, F439I, P440L, and F439I/P440L respectively. Each sample was measured in a series of 4 dilutions to assess intermolecular interactions. The P440L and F439I/P440L mutant peptides aggregated immediately upon dilution into assembly buffer, producing visible macroscopic aggregates; therefore, higher urea concentrations were used prior to measurement. SAXS data were collected at beamline BM29, ESRF, Grenoble, France, using an automatic robotic sample exchanger, measured at room temperature (25 C °).

**SAXS Data Reduction and Analysis**

To determine the inter-filament spacing, two-dimensional diffraction scattering data were collected from neurofilament hydrogels housed in quartz capillaries. Data were azimuthally integrated, and intensity was plotted against the reciprocal distance (*q*). A baseline background equation:

$$I_{bk}(q) = A_0 q^{-\gamma} + C \qquad \text{(S3)}$$

Was subtracted to evaluate the correlation peaks using an inverse Gaussian function(s):

$$I_x(q) = A_1 e^{\frac{-B_1(q-q_1^*)^2}{q}} + A_2 e^{\frac{-B_2(q-q_2^*)^2}{q}} + A_0 q^{-\gamma} + C \qquad \text{(S4)}$$

Here, $A_{0,1,2}$, $B_{1,2}$, C, and γ are fitting parameters. Following, the average interfilament distances were extracted from the peak positions:

$$D_{1,2} = \frac{2\pi}{q_{1,2}^*} \qquad \text{(S5)}$$

To extract local anisotropy (χ-plot), the 2D data were integrate radially surrounding the correlation peak position ($q$=0.087-0.29 $nm^{-1}$) and plotted against the azimuthal angle (χ). Empirically, we found the χ-plot fit:

$$I_x(\chi) = I_{iso} + A_\chi cos\,(2\chi - 2\varphi_D) \qquad \text{(S6)}$$

where $A_\chi$ represents the anisotropy amplitude, $\varphi_D$ indicates the preferred director orientation in real space, and $I_{iso}$ is the isotropic background.

For the peptide, both WT and F439I samples showed a typical IDP scattering pattern that could be fitted to a polymer with a nearly concentration-independent SAXS profile. Therefore, concentrated normalized scattering curves were globally fitted to the extended Guinier formula[4]. The analysis results in the radius of gyration ($R_g$), zero intensity ($I_0$), and scaling factor ($\nu$).

For P440L, which showed a bell-shaped Kratky plot and aggregation in the absence of urea, the SAXS intensity was extrapolated by fitting the low q-range to:

$$I(q \to 0) = I_0^c ex(-B_1 q^2 - B_2 q^4) \qquad \text{(S7)}$$

Where $B_{1,2}$ are fitting parameters and $c$ is the peptide concentration (see supplementary Fig. S12). For a non-aggregating polymer $I(q \to 0) \propto c$ and therefore $\frac{I_0^c}{c}$ is approximately constant. However, for aggregating polymer such as P440L and F439I/P440L, $\frac{I_0^c}{c}$ is monotonically increasing with increasing peptide concentration, and for each urea concentration, we linearly extrapolated $I_0^0/c(0)$ at zero peptide concentration (c→0) and zero SAXS angle (q→0).

The average number of peptides in P440L and F439I/P440L aggregates (i.e., aggregation number, $Z_{agr}$) for a given urea concentration was estimated by:

$$Z_{agr} = \frac{I_0^0(P440L)/c(0)}{I_0^0(WT)/c(0)} \quad \text{(S8)}$$

These values are summarized in supplementary table S1.

**Hydration/dehydration experiment**

Samples for the hydration/dehydration experiment were all prepared and weighed in parallel, with the samples kept in a desiccator between weighing to maintain a controlled humidity environment. The desiccator did not contain any desiccant and remained in room temperature. For rehydration, heated water (50C◦) was placed inside the desiccator. The samples were placed in a 3D-printed holder designed to match the hydrogel size, with a glass bottom to allow observation of the hydrogel under the CPM microscope.

**Bioinformatic and Deep-Learning Ensemble Predictions**

Intrinsic disorder score across the NFL C-terminal tail domain was predicted using PONDR-VLXT[5], Metapredict[6], and IUPred3[7], which estimate disorder propensity from amino acid composition and sequence context. Sequence-dependent IDR ensemble contact maps were generated with STARLING[8] v1.0.0.dev, the software was configured to sample 500 conformations with 40 denoising steps. All predictions were performed locally on an NVIDIA A100 GPU within a Conda-managed environment.

**Statistics**

Statistical analyses were conducted using GraphPad Prism. For capillary SAXS, the sample size N corresponds to the number of independent SAXS measurement per capillary at distinct positions. Dunnett's multiple-comparisons test was used to compare each mutant osmolyte-free interfilament distance ($D_0$) versus WT at ($C_s$=160 mM) (Fig.4d).

**Adjusted p-values and effect sizes (WT vs. mutant, Cs = 160 mM; Dunnett-adjusted):**

F402I: mean diff = 2.787, 95% CI [−0.9535, 6.528], p = 0.2247 (ns).

E396K: mean diff = −2.834, 95% CI [−6.575, 0.9065], p = 0.2108 (ns).

F439I: mean diff = 5.581, 95% CI [1.840, 9.322], p = 0.0009 (***).

P470L: mean diff = 15.32, 95% CI [11.58, 19.06], p < 0.0001 (****).

P440L: mean diff = 16.16, 95% CI [12.42, 19.90], p < 0.0001 (****).

F439I/P440L: mean diff = 15.99, 95% CI [12.25, 19.73], p < 0.0001 (****).

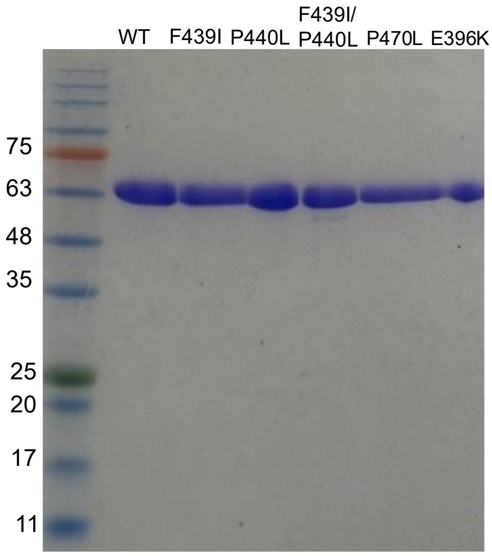


Figure S1. SDS-PAGE analysis of NFL WT and mutant proteins showing purity above 95%.

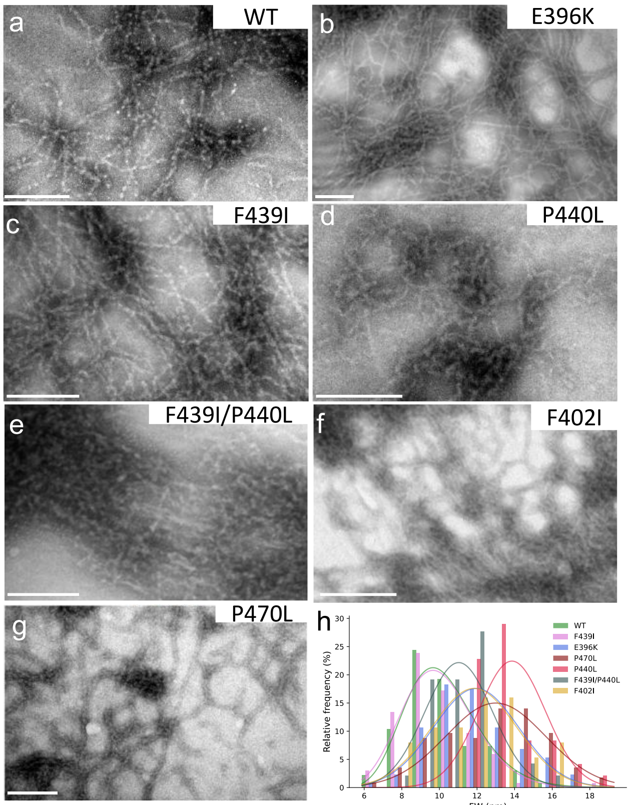


Figure S2. CMT mutations do not disrupt NFL self-assembly**.** TEM images of NFL and mutant samples. **(a)** WT NFL network. **(b-e)** CMT associated mutants P440L, F439I, E396K, and F439I/P440L mutation, respectively. **(f, g)** Control mutation F402I and P470L. Scale bars, 200 nm. **(h)** Filament diameters measured from TEM cross-sections (ImageJ) were similar across variants.

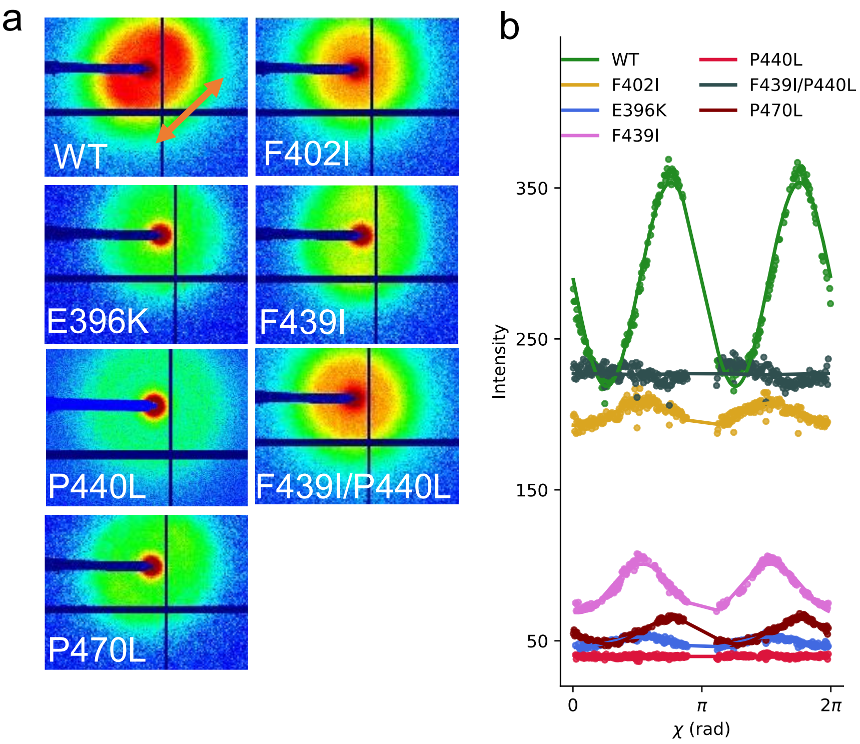


Figure S3. Filament orientation is characterized by SAXS. **(a)** Two-dimensional SAXS scattering patterns of WT and mutant hydrogels. WT displayed the highest scattering anisotropy and the most pronounced directional alignment (i.e., filaments preferentially oriented in a specific direction). **(b)** local anisotropy ($\chi$-plot), was quantified by radial integration around the correlation peak (q = 0.087–0.29 $nm^{-1}$) as a function of the azimuthal angle ($\chi$). WT showed the strongest directional alignment consistent with long-range nematic order, while mutants exhibited reduced anisotropy. The corresponding 1D SAXS profiles are shown in Supplementary Figure S11.

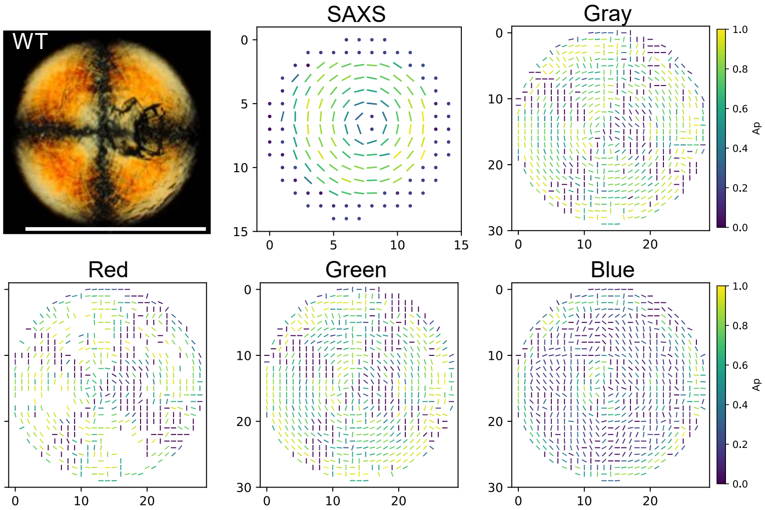
WT
SAXS
Gray
Red
Green
Blue
Ap
0
5
10
15
20
30
0.0
0.2
0.4
0.6
0.8
1.0

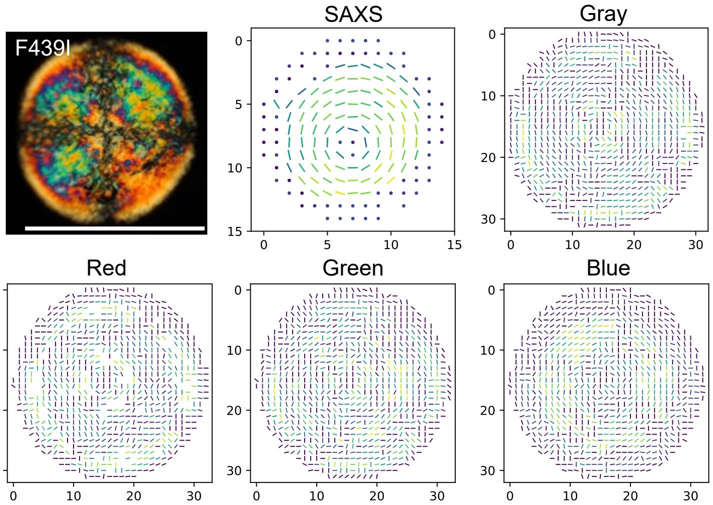
F439I
SAXS
Gray
Red
Green
Blue
0
5
10
15
20
30

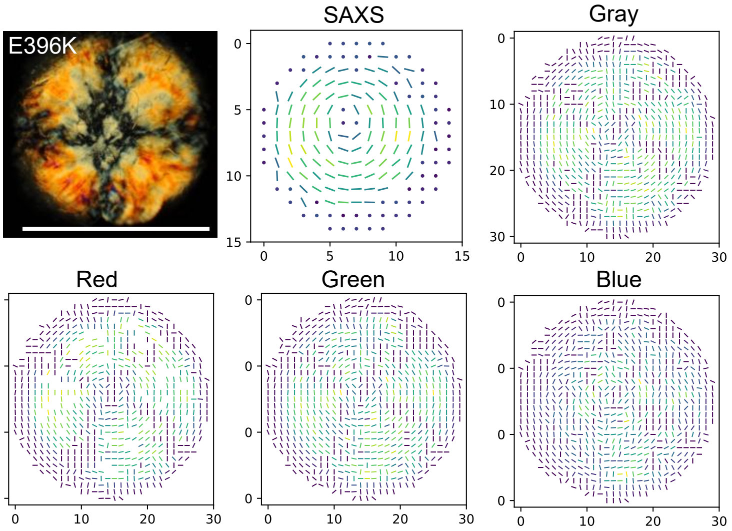
E396K
SAXS
Gray
Red
Green
Blue
0
5
10
15
20
30

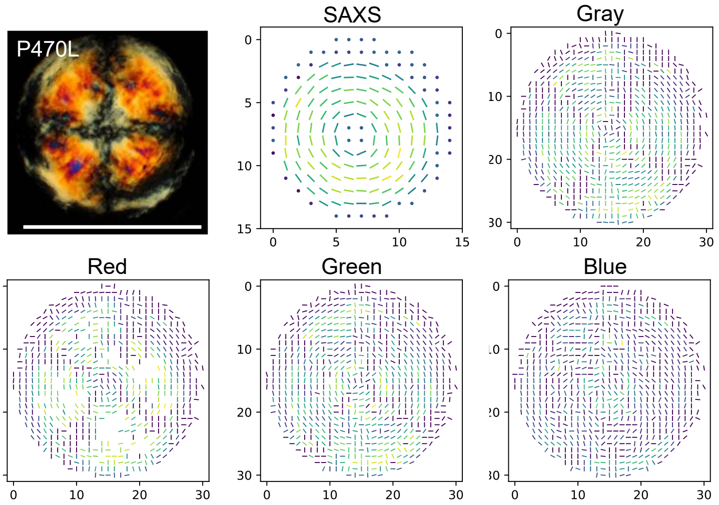
P470L
SAXS
Gray
Red
Green
Blue

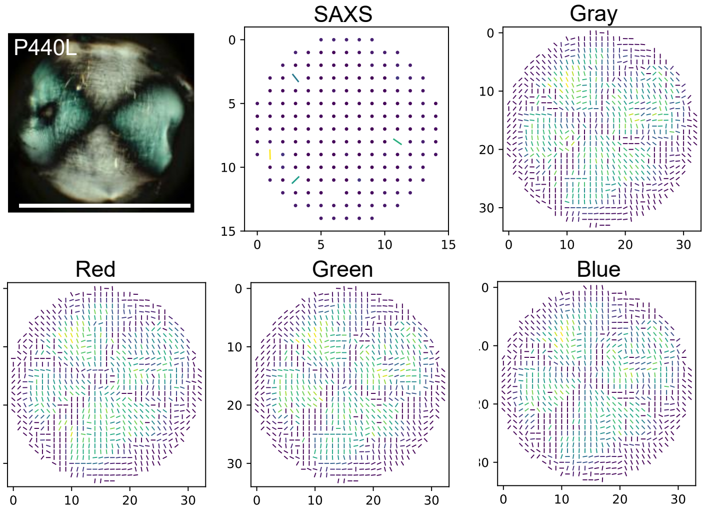
P440L
SAXS
Gray
Red
Green
Blue

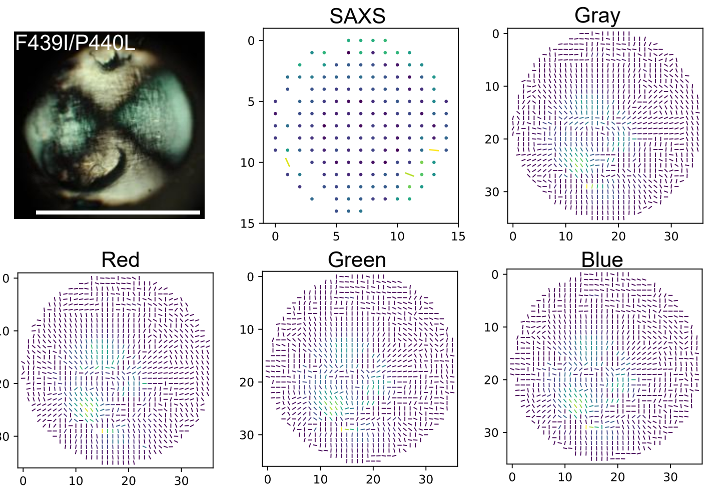
F439I/P440L
SAXS
Gray
Red
Green
Blue

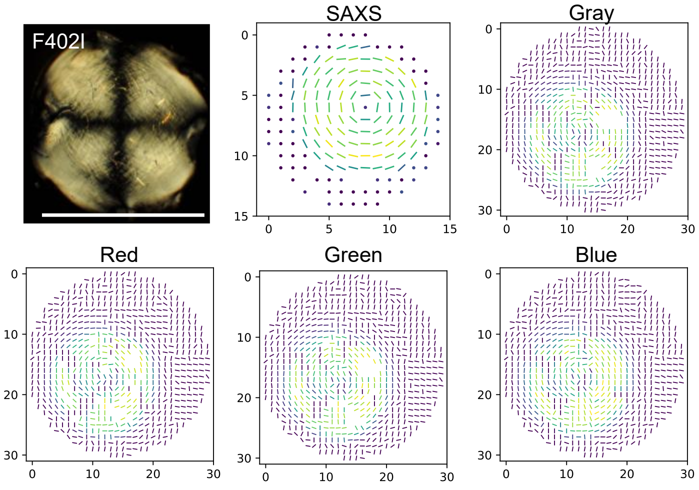


Figure S4. SAXS and CPM mapping of nematic director orientation. Each section shows CPM images, SAXS-derived nematic director orientation maps, and CPM filament orientation maps (grayscale and RGB channels) for NFL hydrogels (Cs = 160 mM, pH ≈ 6.8). In areas where scattering appeared isotropic, the local director orientation map is shown as a point. CPM images reveal two distinct organizational phases: extended nematic ($N_G$) and nematic microdomain ($N_{\mu D}$). SAXS diffraction analysis shows that WT, F439I, E396K, F402I, and P470L hydrogels exhibited a central bend-dominated +1 topological defect with planar anchoring at the periphery. In contrast, P440L and F439I/P440L mutants showed little to no anisotropy in SAXS, likely because domain sizes were smaller than the X-ray beam (~200 × 90 µm²). CPM filament diffraction mapping, corrected for the intrinsic π/2 phase ambiguity using SAXS, showed that WT, P440L, F439I/P440L, and F402I hydrogels had homogeneous birefringence, with WT displaying the highest amplitude. The F439I mutant exhibited colorful microdomains and marked variation in local director orientation across color channels. E396K and P470L mutants displayed intermediate behavior, particularly in the blue channel. scale bar 7.5mm.

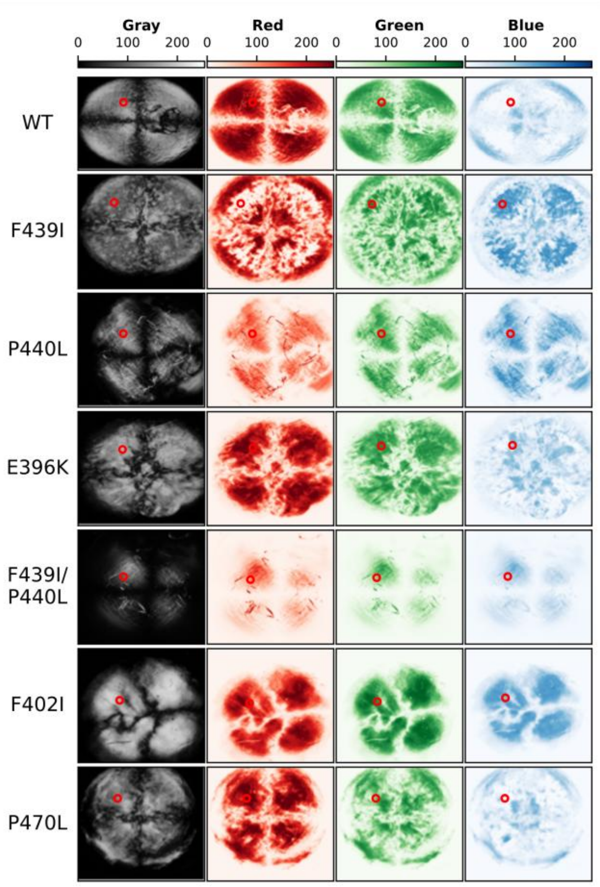


Figure S5. CPM images across color channels**.** Each row corresponds to a different NFL sample (WT, F439I, P440L, E396K, F439I/P440L, F402I, P470L), and each column shows the CPM image for (grayscale, RGB channels). The red point marks regions of intensity quantification (see Supplementary Figure 6) Color bars indicate the intensity scale for each channel.

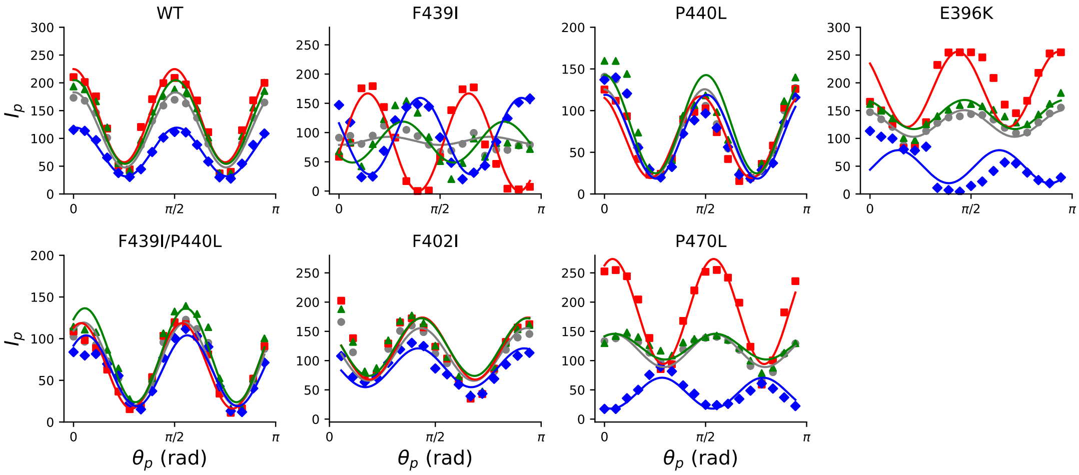


Figure S6. Channel-dependent intensity profiles of NFL hydrogels. Representative intensity profiles were measured as a function of the angle between the sample and the polarizer ($\theta_p$) for (grayscale, RGB channels). WT displayed consistent maxima across all color channels. P440L, F439I/P440L, and F402I mutants showed similar uniform behavior. In contrast, F439I displayed a pronounced channel-dependent shift in maxima, reflecting nematic microdomains. E396K and P470L mutants showed milder heterogeneity.

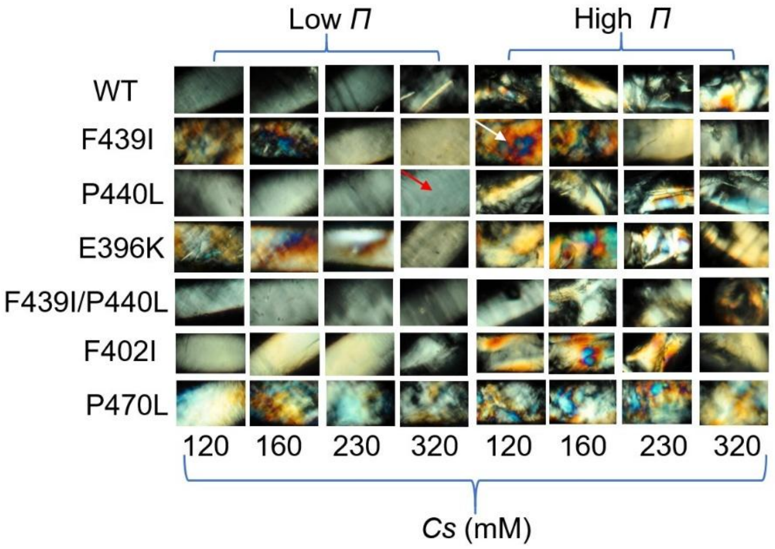


Figure S7. Phase behavior under varying ionic strength and osmotic pressure. CPM images of NFL protein hydrogels in quartz capillaries (~1.5 mm width) at different ionic strengths ($C_s$ = 120, 160, 230, and 320 mM) and varying $\Pi$. Two distinct phases were observed: $N_G$ phase (uniform birefringence; red arrow) and $N_{\mu D}$ (reddish/blue heterogeneous domains; white arrow), characterized by macroscopic reddish/bluish domains. At high $\Pi$, all samples exhibit the $N_{\mu D}$. At low $\Pi$, WT, P440L, F439I/P440L, and F402I retained the $N_G$ phase across all ionic strengths, whereas F439I, E396K, and P470L formed $N_{\mu D}$.

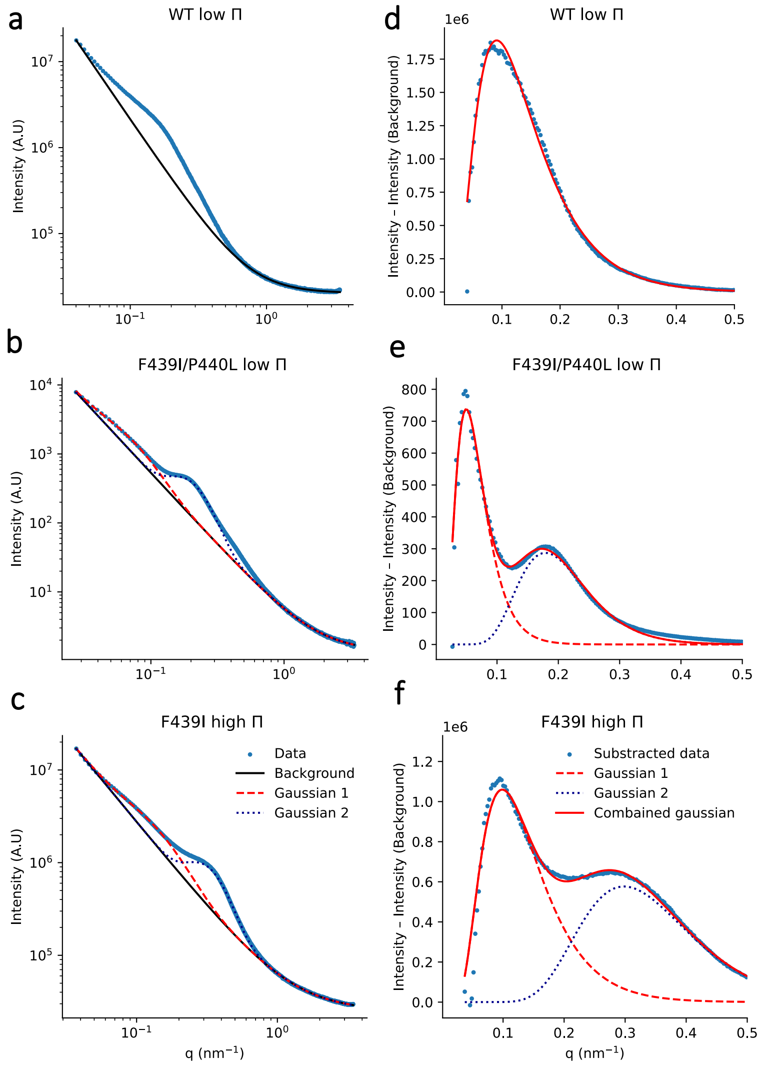


Figure S8. SAXS quantification of interfilament spacing. Representative SAXS intensity curves on NFL hydrogel raw data **(a-c)** and corresponding baseline-subtracted data (**d-f). (a)**WT at low Π (0% w/w PEG), (**b)** F439I/P440L double mutant at low Π (0% w/w PEG), and **(c)** F439I mutant at high Π (≈25% w/w PEG). Broad peaks with single or double maxima were observed. Background scattering (black solid line) was subtracted from the raw data for analysis. **(d-f)** The background-subtracted profile was fitted to an inverse Gaussian function (dotted and dashed lines) to extract the average interfilament distances from the peak position.

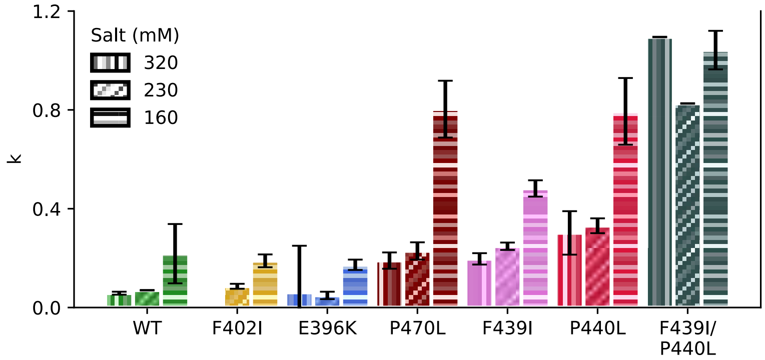


Figure S9. Mean-field compression model of network rigidity. Comparison of the fitted parameter ($k$) from the mean-field model of hydrogel compression theory[2] at varying ionic strengths ($C_s$ = 160, 230, and 320 mM). For larger k values, the network is denser and more rigid at low $\Pi$ with smaller interfilament distances, as shown for the CMT disease-induced mutations: P440L, F439I, and the F439I/P440L double mutation.

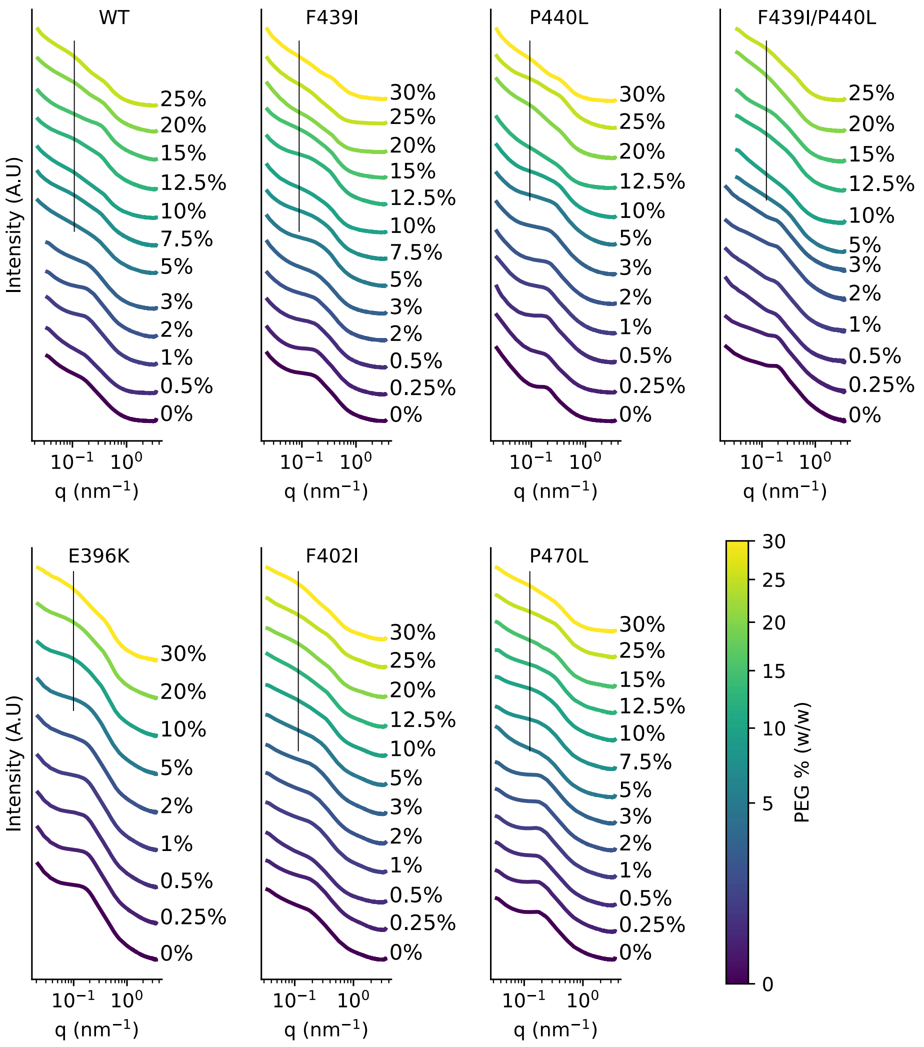


Figure S10. Indication for secondary SAXS correlation peaks. SAXS intensity curve for different samples at varying PEG concentration, labeled next to the corresponding curve. Under high Π (≈5% w/w PEG), all samples displayed a distinct secondary scattering peak at low q values (marked with a black line). The exact position of the secondary peak showed no clear trend with Π, and its q-position remained consistently low (large D values). All measurements shown here were conducted at physiological salinity (Cs = 160 mM), but similar behavior was observed at higher salinities (Cs = 230, 320 mM).

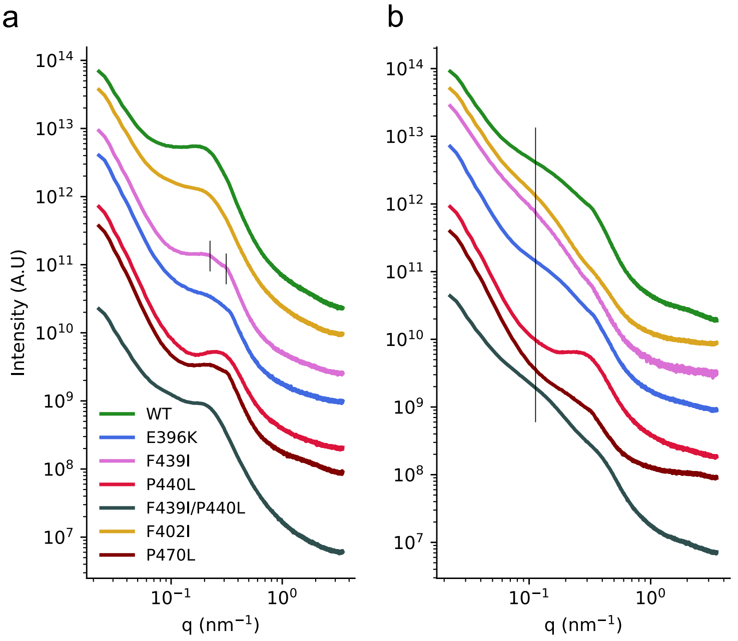


Figure S11. Nematic order and compaction in large hydrogels. **(a)** SAXS intensity curve for large NFL hydrogels. WT showed peaks at smaller q values (corresponding to larger interfilament spacing). F439I, E396K, and P470L samples showing NμD under CPM, exhibited two closely spaced peaks (marked with a black line). The correlation peaks for each sample appear at higher q values than in supplementary figure 10, most likely due to potential air contact or minor dehydration with the scanning SAXS sample holder. **(b)** After exchanging buffer to 25% w/w PEG (high Π condition), the hydrogels exhibited two correlation peaks, indicating the coexistence of distinct interfilament spacing. Secondary peaks are marked with a black line**.**

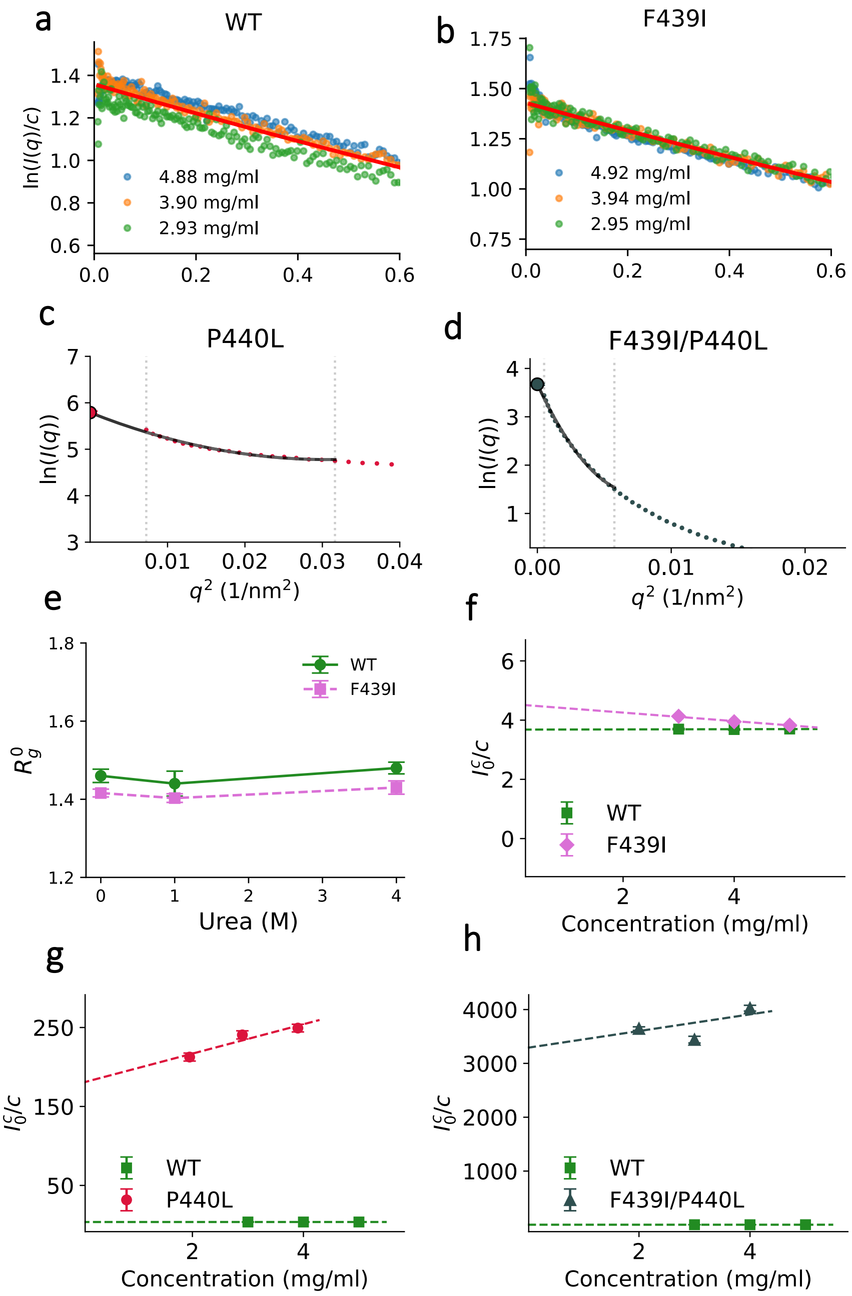


Figure S12. Peptide solution SAXS reveals local structural differences. **(a)** WT and **(b)** F439I peptides show polymeric structural characteristics that can be fitted using the Extended Guinier analysis[4]. **(c, d)** F439I/P440L and P440L show aggregation behavior with access scattering at low $q$. Here, $q$→0 intensity ($I_0$) was estimated by fitting the low $q$-range to $I_x(q) = I_0^c exp\,(-B_1 q^2 - B_2 q^4)$ where $B_{1,2}$ are fitting parameters and C is the peptide concentration. **(e)**WT and F439I showed nearly independence of $R_g$ to C or urea concentration. **(f-h)** $\frac{I_0^c}{c}$ as a function of peptide concentration, values are approximately constant for WT and F439I, whereas for F439I/P440L and P440L, $\frac{I_0^c}{c}$ increases monotonically with C.

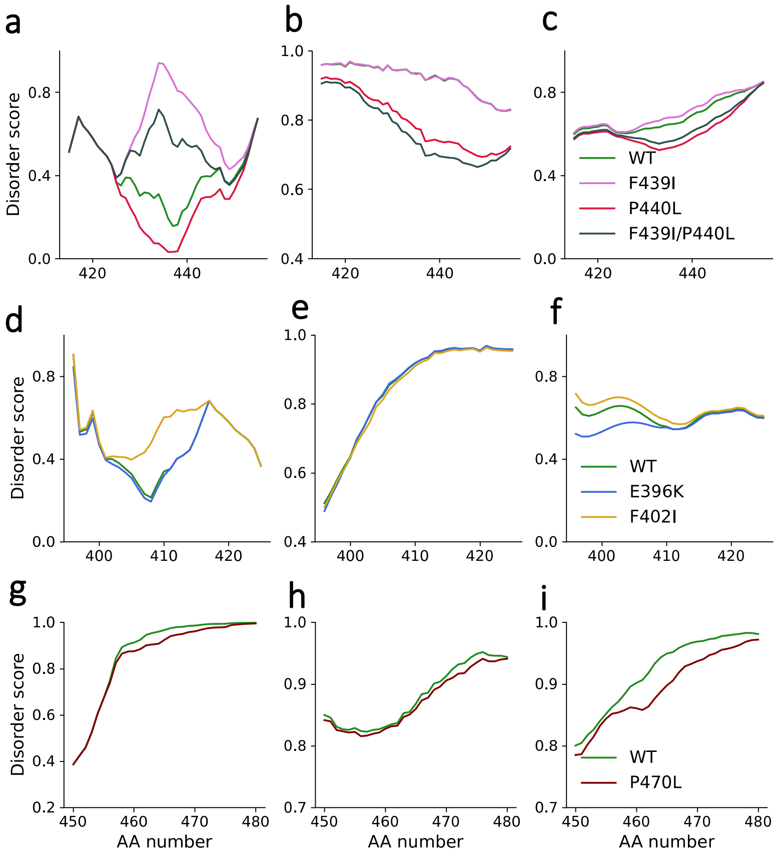


Figure S13. Sequence-based disorder prediction. Disorder scores across the NFL C-terminal tail domain predicted with **(a-g)** PONDR-VLXT[5], **(b-h)** Meta predict[6], and **(c-i)** IUPred3[7]. These tools assess intrinsic disorder based on amino acid composition and sequence context. F439I increased predicted disorder relative to WT, whereas P440L showed a shift toward higher order. Predictions were consistent in trend across tools, though magnitudes varied.

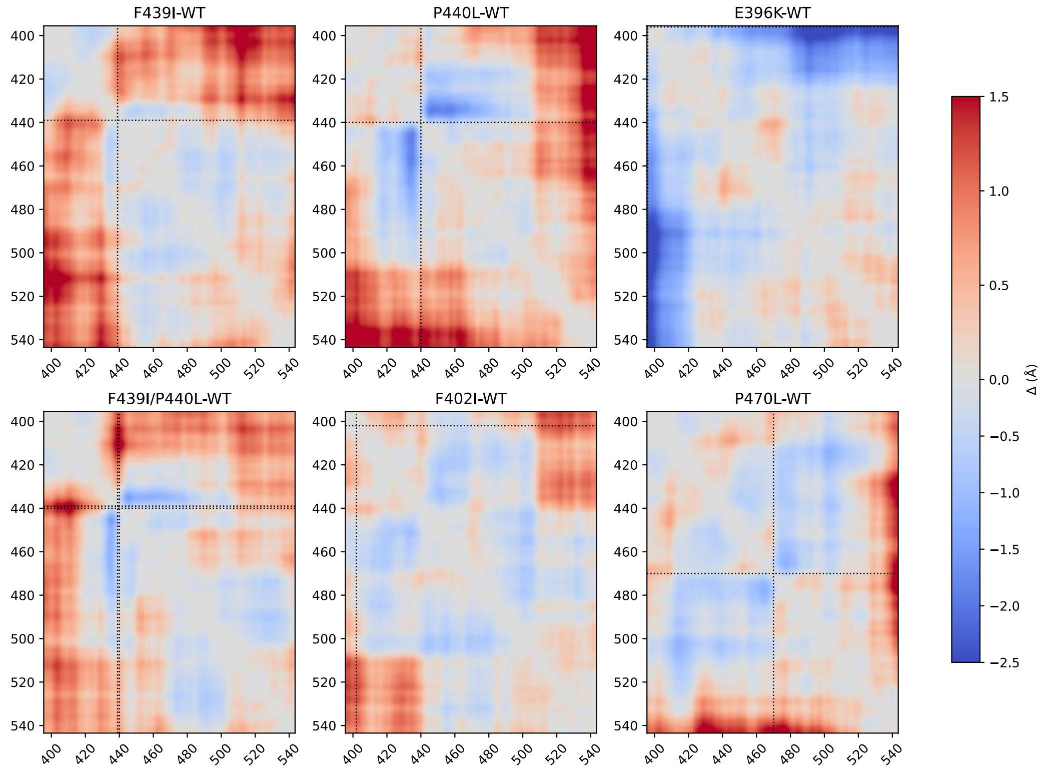


Figure S14. Deep-learning predictions of IDR ensembles' distance maps differences. Deep-learning STARLING[8] prediction of residue-residue distance maps subtracted from the WT prediction (Δ) of the full NFL tail distance map difference from WT P440L, F439I/P440L, and P470L mutations resulted in reduced average inter-residue distances, compared to the WT, in the vicinity of the mutation. In contrast, the F439I and the F402I mutations mildly increased the inter-residue distances. E396K shows decreased average inter-residue distances for full overlap and increased average inter-residue distances for the tip-to-tip contacts. Dashed lines mark the mutation position.

| | Urea conc. (M) | Method | $I_0^0$/ C (0) | $R_g^0$ (nm) | $\nu$ | $Z_{agr}$ |
|---|---|---|---|---|---|---|
| **WT** | 0 | Extended Guinier | 3.21±0.18 | 1.46±0.02 | 0.604±0.001 | 1 |
| | 1 | Extended Guinier | 3.7±0.04 | 1.44±0.03 | 0.606±0.001 | 1 |
| | 4 | Extended Guinier | 2.7±0.003 | 1.48±0.015 | 0.607±0.002 | 1 |
| **F439I** | 0 | Extended Guinier | 4.29±0.09 | 1.42±0.01 | 0.606±0.002 | 1 |
| | 1 | Extended Guinier | 4.54±0.09 | 1.403±0.01 | 0.576±0.002 | 1 |
| | 4 | Extended Guinier | 3.10±0.32 | 1.43±0.02 | 0.612±0.003 | 1 |
| **P440L** | 0 | Extrapolation | 29.65±1.20 | — | — | 9.23±0.64 |
| | 1 | Extrapolation | 179.74±17.91 | — | — | 48.58±4.87 |
| | 2 | Extrapolation | 416.03±8.26 | — | — | 274.42±7.94 |
| **F439I/ P440L** | 0 | Extrapolation | 2629.66±1106.4 | — | — | 819.6±347.7 |
| | 1 | Extrapolation | 7971.34±13.33 | — | — | 2154.4±23.6 |

Table S1. SAXS analysis of NFL peptides under varying urea concentrations. Extrapolated peptide concentration normalized to zero-concentration scattering intensity $I_0^0$/ C (0) and zero-concentration scattering radius of gyration $R_g^0$. Aggregation number ($Z_{agr}$) of P440L as described in the methods.
